# QT-RelCor: Statistical Inference for Comorbidity of Quantitative Traits and Disease from Related Individuals

**DOI:** 10.64898/2026.11.16.766965

**Authors:** Prasun Panja, Pavithra Jayasankar, Swarna Buddha Nayok, Biju Viswanath, Meera Purushottam, Sanjeev Jain, Samsiddhi Bhattacharjee

## Abstract

**Background:** Complex diseases with a high genetic component often cluster in families and are associated with quantitative traits (QT-s) such as endophenotypes showing comorbidity with the end-point disease trait. Detecting associations with such QT-s can provide a non-invasive approach to indicate disease onset at an early stage, or help in tracking disease severity. Traditional case-control studies generally involve unrelated individuals, which can introduce heterogeneity due to varying lifestyle and exposure patterns. Often QT values may be available from related individuals such as families of affected probands. In these situations, assessment of comorbidity restricting to unrelated cases and controls leads to unnecessary loss of power. On the other hand, ignoring the corelation structure of families leads to grossly incorrect confidence intervals impacting both type 1 error and power of tests. Yet, these practices are common in the absence of standard inference procedures for such data.

**Methods:** We propose statistical procedures for analyzing clustered data to infer association between the QT-s and the binary trait. The methods allow a mix of families and unrelated cases and utilizes all available individuals. In particular, two novel competing approaches are proposed and evaluated: a parametric approach based on multivariate normal likelihood and a non-parametric approach that combines the Wilcoxon rank-sum test with a modified version of the Wilcoxon signed-rank test.

**Results:** We conducted simulation studies to assess type 1 error and power across multiple scenarios, which demonstrated the robustness of the proposed tests. Both the tests perform considerably better than both the classical t-test and Wilcoxon signed-rank test under both parametric and non-parametric settings. We illustrate the proposed methods by applying them on sibship data from the Centre for Brain and Mind program to identify a number of quantitative precursors significantly associated with schizophrenia and bipolar disorder.

**Conclusion:** In summary, the proposed methods provide a robust and powerful approach to detect comorbid endophenotypes incorporating data from families of affected individuals. Findings from the analysis of sibling-based data on schizophrenia and bipolar disorder support role of a few selected neurocognitive assessments as potential markers of these disorders.

## 1 Introduction

Different health outcomes in an individual may tend to co-occur, due to shared pathogenic mechanisms, or because one condition or its treatment triggers another [1, 2]. For example, in long-standing type-2 diabetes, it is common to have diabetic neuropathy, retinopathy, or nephropathy. Such co-occurrence of various phenotypes in an individual along with the primary phenotype of interest, regardless of any causal relationship, is termed as comorbidity [3]. Identifying which conditions or traits are associated with the primary disease phenotype could inform preventive and therapeutic interventions for multiple conditions collectively [4]. In many cases, due to the presence of age and other potential confounders, it is unclear whether there is a non-random association between two phenotypes [5–7]. For example, coronary artery disease (CAD) often co-occurs with elevated systolic blood pressure (SBP) levels or hypertension. Mechanisms for such comorbidity are not always obvious, underscoring the need for statistical approaches to inferring patterns of co-occurrence between the traits.

Comorbidity is highly prevalent in cases of major psychoses such as schizophrenia or bipolar disorder (BPAD). Several quantitative traits involving various neurosensory activities, reflex actions, or tasks related to auditory and memory functions are suspected to be correlated with such psychoses. Analyzing these quantitative traits to assess their comorbid relationships with binary traits like schizophrenia or BPAD would simplify the process of diagnosing the binary trait at early stages and assist in developing treatment/management guidelines based on these traits. Identifications of endophenotypes can be the foundation for community based tools for screening or monitoring of such binary diseases [8, 9]. This concept applies generally to most chronic diseases that can be identified through various quantitative precursors.

There are two distinct study designs to analyze comorbidity. The first is a population-based approach, which involves recruiting unrelated cases and controls. While this design is widely used [10, 11], it has to account for considerable interindividual variation. For example, Age, sex, ethnicity, and socioeconomic status also influence comorbidity patterns [12]. The second approach involves recruiting diseased individuals and using their unaffected first-degree relatives as controls. Unlike population cohorts, which typically mix diverse backgrounds, family-based data can offer sharper inference by reducing heterogeneity in background risk factors since both genetic and environmental factors are known to contribute to the comorbidity [13]. However, this design is challenging for late-onset diseases, as it can be difficult to recruit controls from previous generations (e.g., parents). A common workaround is to use unaffected siblings as controls [14], but this is not always feasible as some individuals may not have siblings, leading to their exclusion from the study. Hybrid study designs that effectively integrate both population-based and family-based approaches can partly address these limitations. Family-based and hybrid designs are commonly used in genetic epidemiology for studying genomic variants [15–17]. They can improve statistical power and reliability by maximizing the inclusion of eligible participants, including families.

Existing approaches to quantify comorbidity can be broadly divided into two categories. The first category examines the non-random co-occurrence of diseases, often using odds ratios (ORs) or relative risks (RRs) within classical case-control designs. Such approaches have been applied, for example, to study the link between pyrethroid insecticide exposure and ADHD [10], and to assess the impact of neighborhood poverty on heart failure incidence [18]. While informative for identifying comorbid endophenotypes, these designs remain limited to pairs of binary or categorical phenotypes and are susceptible to population heterogeneity and confounding. Beyond these, network-based methods such as Morbinet[19] leverage large-scale medical records to map multimorbidity patterns and disease trajectories [4, 20–22]. However, hospital based data often miss common chronic conditions that are typically managed outside hospital settings. Moreover, their primary focus is on disease progression networks rather than identifying trait-level precursors. Although effective in summarizing large cohorts, these methods remain largely descriptive and lack the ability to formally test comorbidity hypotheses.

In this study, we focus on identification of quantitative traits (not always classified as diseases) that may inform the diagnosis of binary endpoint diseases or provide insights into disease manifestation. This is a commonly encountered situation that has been received relatively less attention. Previously, association of such traits have been explored using group comparisons between patients, their first-degree relatives and healthy individuals primarily using ANOVA and ANCOVA models of the quantitative traits on disease outcomes such as bipolar disorder or schizophrenia [14, 23] or other binary traits [24, 25] adjusting for demographic and other covariates. Another approach utilizes the reverse model of disease (Alzheimer’s) on a quantitative trait (clock-drawing errors) using a logistic regression [8]. Approaches other than regression have also been attempted. A study employed EM-based finite mixture modeling to examine whether the distributions of schizophrenia endophenotypes such as sustained attention deficit, and eye-tracking dysfunction in unrelated individuals have latent multi-modal structure [26].

While these studies provided useful insights, application of these analytic methods for correlated individuals such as first-degree relatives of affected individuals without accounting the correlations might lead to inflated false-positive rates. Restricting analysis to a subset of independent individuals would lead to substantial power loss. Further, the methods discussed are based on parametric assumptions, and hence can not be expected to work optimally for non-normal or skewed traits. Mis-specification of the trait distribution may lead to reduced statistical power and inflated type I error rates. Dobri et. al [14] used a non-parametric method, i.e., Kruskal-Wallis test to allow for non-normal traits. However, this test is limited to one-way ANOVA only and is also not suitable for related individuals. These limitations motivate the need for developing statistically rigorous frameworks, particularly those suited for correlated data structures such as families.

In this article, we propose statistical procedures to study the association of a quantitative trait with a binary trait (usually disease) using data on independent patients as well as sibships comprising both probands and related controls. Specifically, we propose two complementary strategies: one that operates under a parametric framework assuming normality, and another that is non-parametric, making no strong distributional assumptions. Since our methods rely on the comparisons with the controls from the same sibship, the potential effects on variations in environmental risk factors such as lifestyle, dietary habits, or socioeconomic background are minimized (see section 2). Moreover, the probands and their paired controls are from the same generation of the same family further reducing variability. Our methods can also incorporate the population-based case-control data when available and combine with the family-based data to derive maximal power while correctly accounting for the relatedness.

The statistical methods developed here can potentially be applied more generally in other contexts such as in analysis of molecular quantitative traits. This could include cancer gene-expression data from a mix of paired and unpaired tumor and adjacentnormal samples or even data from single-cell gene-expression studies, where cells within an individual comprise a correlated cluster. We evaluate the statistical properties of these methods through comprehensive simulation studies, and illustrate their application using data from the Centre for Brain and Mind cohort, focusing on schizophrenia and bipolar disorder and several quantitative neurocognitive assessments.

## 2 Comparison of Study Designs

It is arguable whether a family-based design provides any advantage over the standard case-control approach for identification of comorbid or correlated traits. In this section we argue that indeed, the family based design (represented by paired data) can be substantially more powerful than unrelated cases and controls (unpaired data). Intuitively, this is because, pairs are highly correlated when the noise (i.e. unmodeled factors) are similar within the pairs.

To support this argument, we conducted a power comparison between the paired t-test and the two-sample t-test with a sample size of 50 and an error variance of 2. The results are summarized in Figure 1. We observe that, under similar settings, the power of the paired t-test not only increases with the correlation coefficient but also consistently exceeds that of the two-sample t-test across all levels of correlation, particular for small effect sizes. The methods and simulation setting for the comparison are detailed in the Supplementary Methods section S1.1

**Fig 1:**
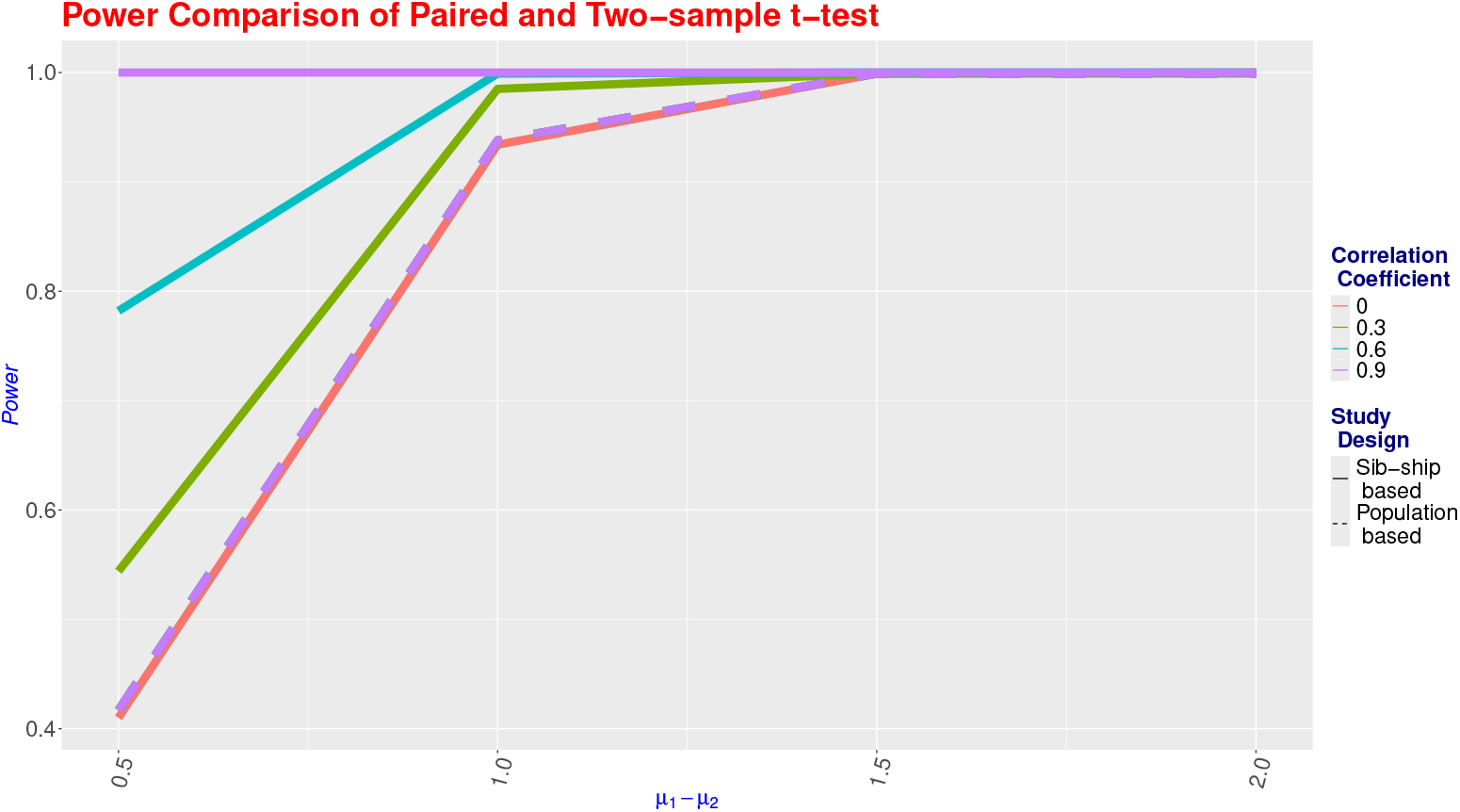
Theoretical power comparison of paired and unpaired t-tests with respect to correlation coefficient and effect size

## 3 Methods

Here, we consider a general framework that can be used to estimate associations based on datasets consisting of groups of correlated observations, which we refer to as ‘families’ or ‘sibship’. The data comprises case individuals (e.g. those affected by a disease), for some of whom additional related family members (e.g. siblings) are recruited. Further, we assume that the families are not necessarily sampled from the population. They could be recruited (ascertained) based on at least one family member being a case. QTs are then measured on all available individuals. The hypotheses therefore relate to marginal differences in QT means between disease groups and the control groups. The data are collected on families that are potentially recruited based on the disease status of probands, while the test procedures account for within-family dependence, and use all available data instead of relying only on family-specific contrasts such as paired tests.

We assume *X*_*ik*_ be the continuous trait of the i-th individual of the k-th family.

*m*_*k*_ be the dimension (i.e., number of family members) of the k-th family. *D*_*ik*_ be the disease indicator such that *D*_*ik*_ = 1 if the i-th individual of the k-th family.

### 3.1 Parametric Setup

Now under parametric setup we assume that

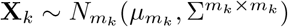

*µ* be the mean vector with each element of the form [*D*_*ik*_*µ*_1_ + (1 − *D*_*ik*_)*µ*_2_] Σ be the dispersion matrix of the form 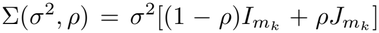, where I is the identity matrix and J is the matrix with all elements is 1. We want to test

*H*_0_ : *µ*_1_ = *µ*_2_ against *H*_1_ : *µ*_1_*≠ µ*_2_.

Under *H*_0_, *µ*_1_ = *µ*_2_ = *µ*. The likelihood of the k-th family (dimension is *m*_*k*_) is,

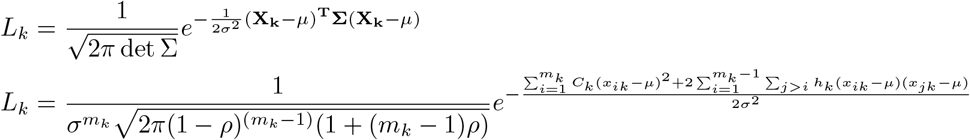

So the likelihood of the data is 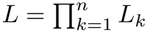 and the log-likelihood *l*(*µ, σ*^2^, *ρ*) is,

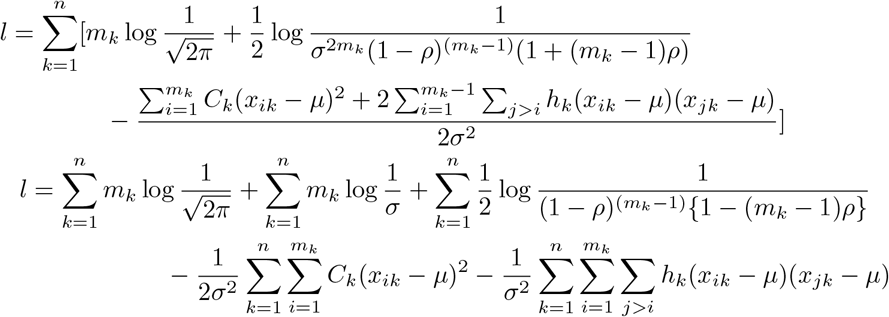

where,

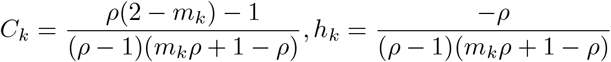

where these are the diagonal and off-diagonal elements of the block correlation matrix corresponding to Σ(*σ*^2^, *ρ*).

we get the likelihood equations by differentiating the log-likelihood w.r.t.the parameters.

To estimate *µ*, we have 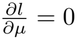, where

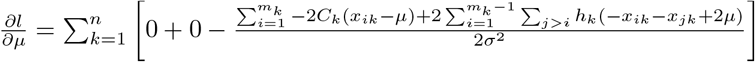

Now we have,

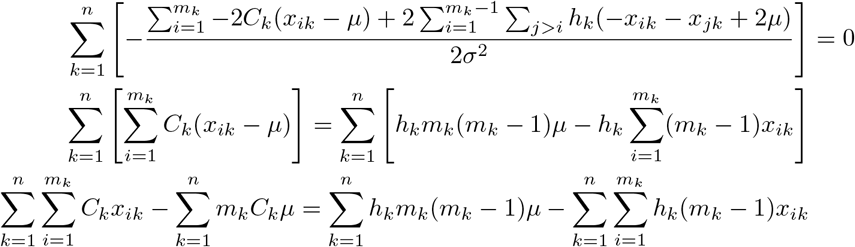

The closed-form profiled mle of *µ* is given by

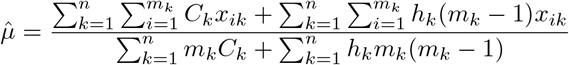

Similarly, to obtain the estimate of *σ*^2^ we have 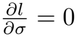, where

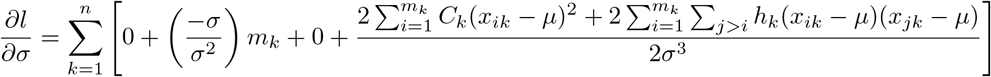

So we get,

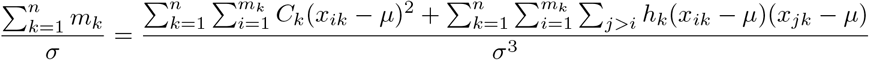

Hence we get the closed form profiled mle of *σ* as

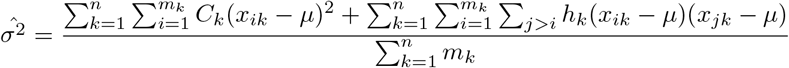

Now we define, 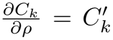 and 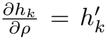 . To obtain the estimate of *ρ*, we have 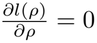, where the *l*(*ρ*) is the profile log-likelihood of *ρ*. Hence, we get

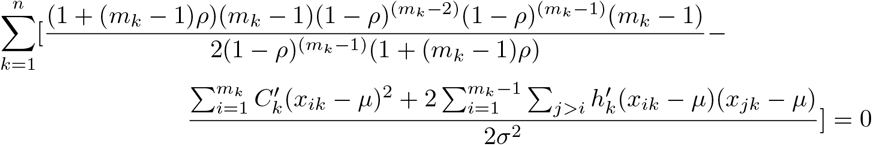

We solve this equation using Newton-Raphson iterations, as it does not have a closed-form solution. We notice that 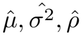 are dependent on each other. So, we start the estimation procedure with an initial guess of *ρ* to obtain the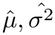 . Then we update *ρ* to obtain again 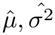 and continue this until convergence.

Under *H*_0_ ⋃ *H*_1_, the log-likelihood *l*(*µ*_1_, *µ*_2_, *σ*^2^, *ρ* is,

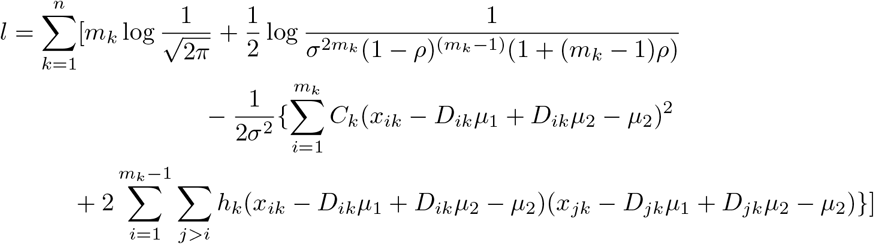

Similarly here we obtain the estimator of all the parameters. For *µ*_1_ and *µ*_2_ we get,

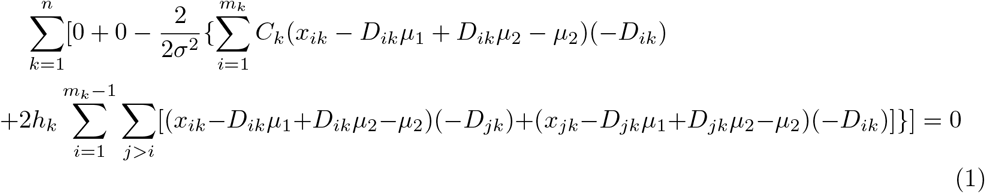

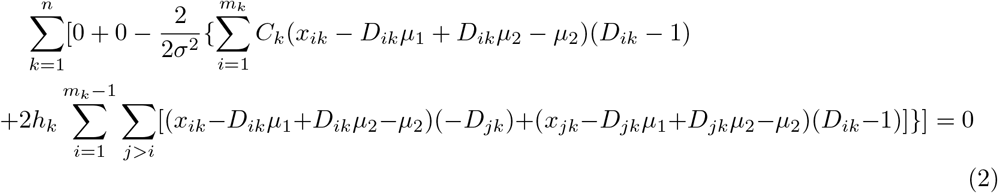

We solve equation (1) and (2) simultaneously to obtain the profiled mles 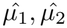. The profiled mles of *σ*^2^ and *ρ* are obtained in the similar way as done under the null distribution.

The likelihood ratio is obtained as 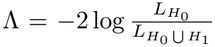, which follows 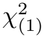 distribution asymptotically. We reject the null hypothesis if 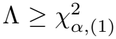 for a given level of significance *α*.

#### Relation to Generalized Least Squares (GLS) and Linear Mixed Models (LMM)

The proposed parametric estimator is equivalent to a Generalized Least Squares (GLS) estimator, when the family covariance structure is known (See Supplementary Section S1.2.1). For example, in terms of the vectors of QT values **X** and binary trait values **D**, these data could be modeled as

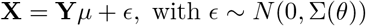

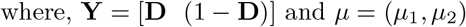

Here we have assumed a compound symmetric covariance model Σ = Σ(*σ*^2^, *ρ*) for each *m*_*k*_ dimensional sibship. We employed iterative estimation of the variance–covariance parameters along with the regression coeffcients under the full ML framework as this is considered to be more efficient than the Feasible GLS (FGLS) approach [27, 28], which uses residuals from an initial OLS model to estimate the variance-covariance matrix.

Another alternative approach is to parametrize the same model under linear mixed model (LMM) framework with a random intercept for each family as

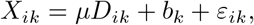

where *b*_*k*_ *∼ N* (0, *τ* ^2^) and *ε*_*ik*_ *∼ N* (0, *ω*^2^). Under this parameterization, the intra-family correlation coefficient (e.g., see Supplementary Methods, Section S1.2.2 and [29]) is *ρ* = *τ* ^2^*/*(*τ* ^2^ + *ω*^2^) = *τ* ^2^*/σ*^2^. Typically for LMM, REML is used to estimate *τ* ^2^ and *ω*^2^. However, when the fixed effects are primarily of interest, full ML estimator (our approach) should be preferred. Wald test of fixed effects under REML is known to show type-1 error inflation and LRT faces problems when the fit becomes singular [30]. However fitting the LMM with REML estimation expectedly gives very similar results as the full ML approach discussed here (see Supplmentary Figure S1).

It is to be noted, that the LMM parametrization is only applicable in the equicorrelation setup. The proposed parametric framework can be readily extended to more complex correlation structures (e.g. see Autoregressive correlation used in Simulations) or families including both parents or extended pedigrees, provided that the structure of the corresponding family covariance matrices are specified. In such cases, the variance–covariance matrix can be expressed as Σ = Σ(*θ*), where *θ* denotes the corrlation parameters estimated iteratively within the Maximum Likelihood (ML) framework using profile likelihood. Further, covariates can be incorporated in a natural way within the GLS framework (see Supplmentary Methods, section S1.3.1).

### 3.2 Non-parametric Setup

To develop nonparametric tests, we make the same assumptions as described above, except that we do not assume the population distribution of the quantitative phenotype to be multivariate normal. The test is divided into two components:

1. The first component is a Wilcoxon rank-sum test for two samples using independent cases and the mean of the family controls only, say *T*_1_.
2. The second component involves a modified Wilcoxon signed-rank test using the mean of family cases and the mean of family controls within the same family, say *T*_2_.

Since the variance of the mean of the family controls varies based on the number of cases and controls in a family, the usual assumption of equality of variances is violated.

Additionally, the variance of the mean of the family controls differs from that of the independent cases. Therefore, we modify the test as follows.

For *m*_*k*_ *>* 1, the mean of the affected sibs of the k-th family is

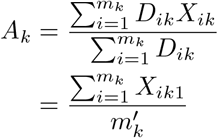

where, *X*_*ik*1_ is the continuous trait value of the affected sib in the k-th family and 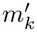 is the number of affected sib in the family. Now we have

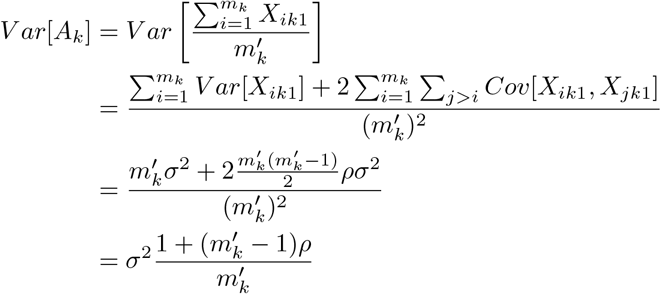

Similarly, the mean of the controls can be written as

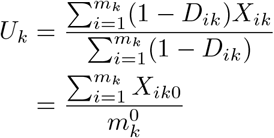

where, *X*_*ik*0_ is the continuous trait value of the unaffected sib in the k-th family and 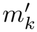 is the number of unaffected sib in the family. The variance of *U*_*k*_ is 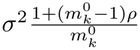 So if we use 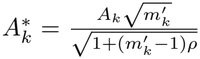 and 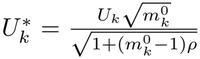 assumption holds. Such scaling can be used to adjust the variance for families with any general structure, as long as the intra-family covariance of the QT is specified. Let _*d*_ denote the set of families with sibship size ℱ *d*. For each sibship-size class, a product–moment intraclass correlation coefficient is computed as

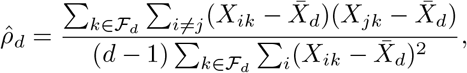

where 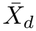 denotes the pooled mean within that class. The correlation used for variance stabilization in the nonparametric procedure is then obtained as a weighted average across sibship sizes,

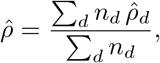

where *n*_*d*_ = | *ℱ*_*d*_ | is the number of families of size *d*. This moment-based pooled ICC serves only as a variance-stabilizing summary of within-family dependence and does not impose a parametric model on the QT distribution.

Let *R*^(*C*)^(*·*) denote the mid-rank function computed after pooling independent cases together with the scaled means 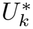 of the family controls. The first component *T*_1_ corresponds to a Wilcoxon rank-sum statistic comparing independent cases with the scaled family controls, and can be written as

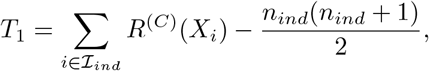

where *ℐ*_*ind*_ denotes the set of independent cases, *n*_*ind*_ is the number of independent cases. Now assume that *R*^(*D*)^(*·*) be the mid-rank function of the absolute differences between the mean of the family cases and mean of the family controls. Let, 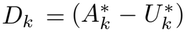 for the *k*-th family and assume that *R*^(*D*)^(*·*) be the mid-rank function of |*D*_*k*_|, if *D*_*k*_ *≠* 0. Now the 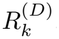, the rank of the absolute differences are signed with the sign of *D*_*k*_. Now the second component *T*_2_, a Wilcoxon signed-rank statistic, can be written as

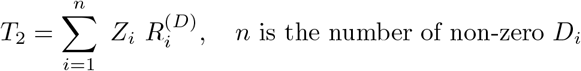

*Z*_*i*_ is the indicator variable such that *Z*_*i*_ = 1 if the difference *D*_*i*_ *>* 0 and *Z*_*i*_ = 0 otherwise. The composite statistic is then *T* = *T*_1_ + *T*_2_.

To asses the extent and direction of association using the proposed nonparametric method for each QT, we compute the difference between the observed test statistic and the expected test statistic under null hypothesis 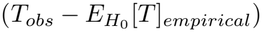. A positive difference indicates a positive association, meaning the presence of the binary trait increases the value of the QT. Otherwise, it is negatively associated. It is worth noting that the variance of the QT also impacts the decision, not just the difference. In fact, as demonstrated in our simulation studies, the power of the proposed test for each QT is an increasing function of the difference in the mean of the QT between cases and controls divided by the standard deviation of the QT (analogous to Cohen’s *d*). The expected value of the test statistic under the null hypothesis and p-values are obtained using a permutation-based algorithm. Specifically, the binary trait labels are shuffled within each family under the null (to preserve the observed dependence structure). Since labels are independent of trait values under the null hypothesis conditional on family membership, this yields valid inference without requiring a parametric covariance model.

#### Algorithm 1

Parametric Likelihood-Based Approach

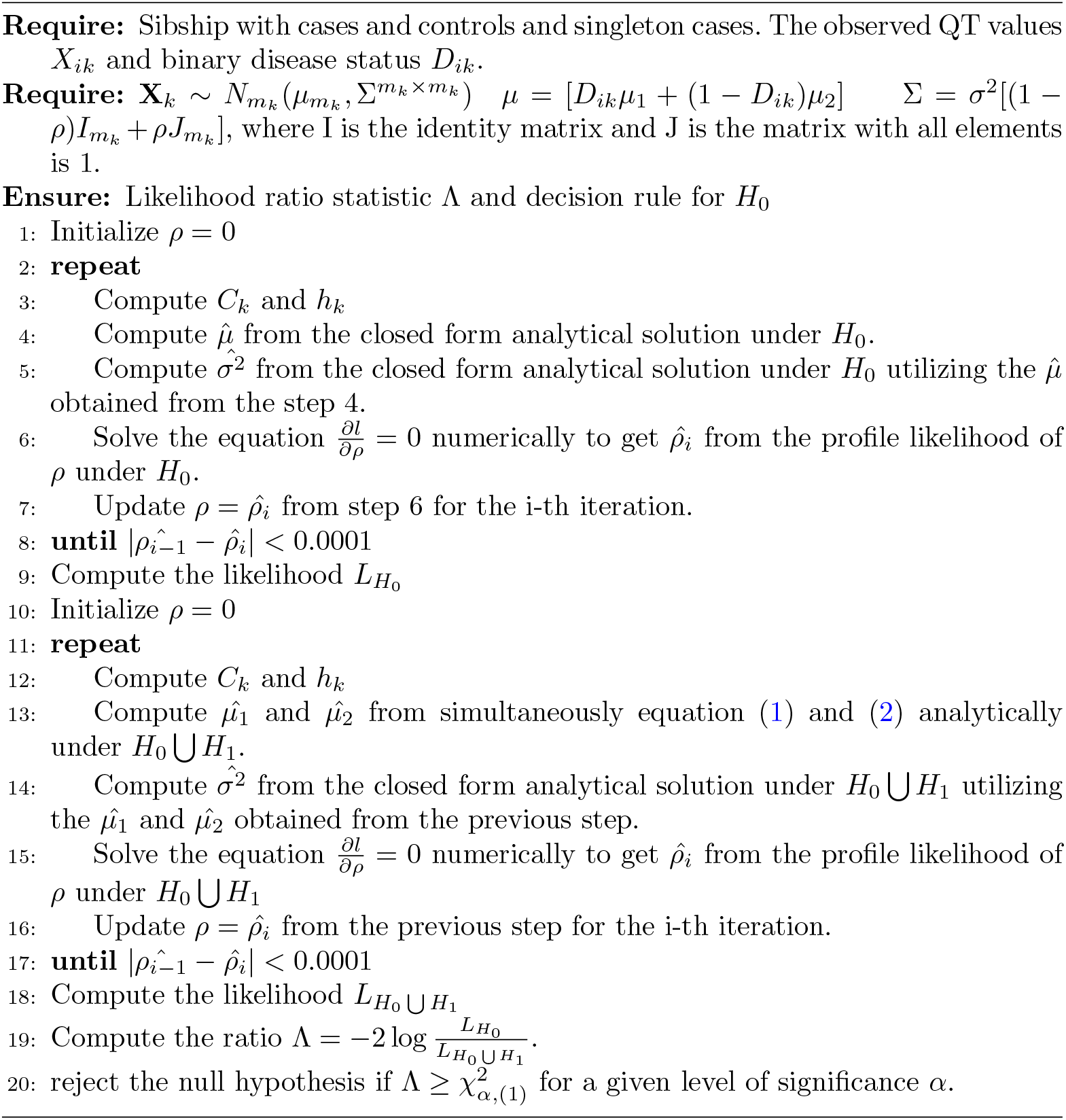

## 4 Simulation Study

We carried out extensive simulations to detect false positive rates and powers for detecting association between the binary trait and its quantitative precursor.

### 4.1 Simulation Models

For simulations, we assumed a model where at least one member of the sib-ship is affected with the binary trait, as otherwise the sib-ship would not provide informative data. Further, assuming that at most 4 siblings can be present in a sib-ship, we generated the number of members in each sib-ship from a truncated Poisson(*λ*) distribution

#### Algorithm 2

Nonparametric Approach

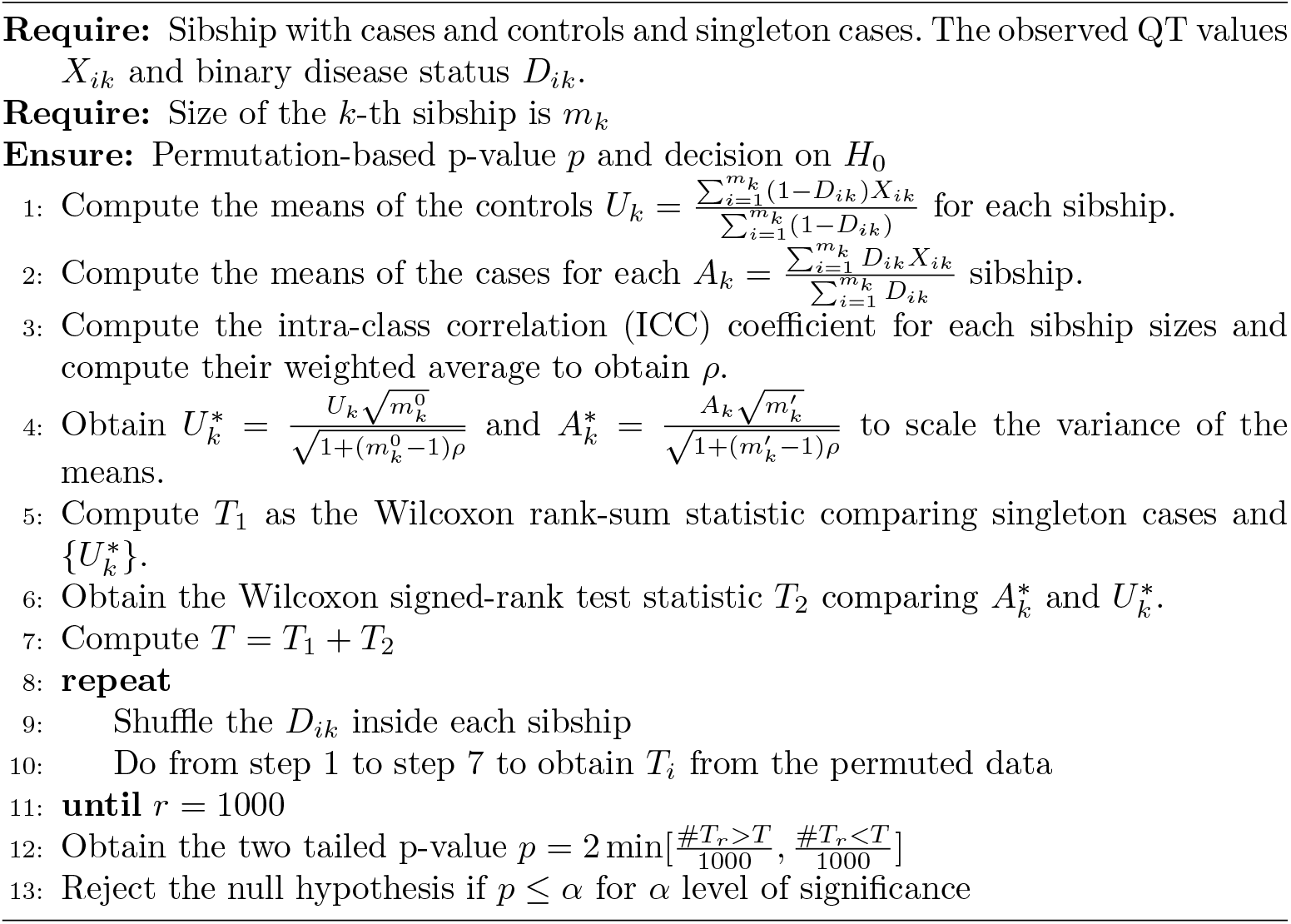

truncated at 0 and above 4. So, if N is the random variable of number of members in a sib-ship, then

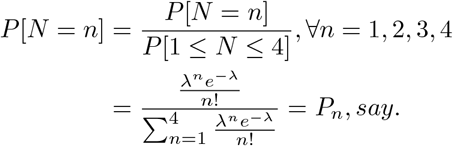

We generated a total *k* number of families from a multinomial(*k,P*_1_, *P*_2_, *P*_3_, *P*_4_) distribution. Let **Y**_**i**_ denote the vector of the binary trait of the i-th family. Obviously, length of the vector is equal to the i-th family size and *Y*_*ij*_ = 1 if the j-th member of the i-th sibship is affected with the binary trait. We generated the number of affected sib in a sibship from a truncated Binomial(*m*_*k*_, *p*) distribution with truncation at 0 and *m*_*k*_, where p is the probability of being affected in the given sibship. *p* is generated from a uniform (0,0.5) distribution. We supposed **X**_**i**_ is the vector of the continuous trait of the i-th family. Here we assumed two types of distribution. The first was **X**_**i**_ follows a multivariate normal distribution with mean (*µ* + *γ*.**Y**_**i**_), where *γ* is a scalar quantity and the dispersion matrix is Σ. The second one was a location shifted multivariate chisquared distribution, where **X**_**i**_ follows a multivariate 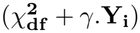 distribution with *Cor*(*X*_*ij*_, *X*_*ij\**_) = *ρ*. When we simulated under null hypothesis *H*_0_, the value of *γ* is set to 0,while under the alternative hypothesis *H*_1_, *γ ≠* 0.

### 4.2 Simulation Settings

We carried out simulations to compare the parametric and non-parametric approach with respect to the false positive rates and powers. We calculated the type I error rates and powers based on 50 families and 1000 replications. For the nonparametric approach, empirical p-values were computed using 1000 permutations. Initially, we set *λ* to 1.2 and *γ* to three different values: 1, 1.5 and 2. To assess the symmetry of the test, we also set *γ* to -1, -1.5 and -2 under both N(0,2) and 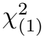 distributions. We also varied the variance of the quantitative traits from 2, 4 and 6 under both normal and scaled 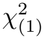 distributions. We set the scale parameter as 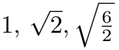. The ratio 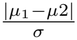 might has the effect on the method and so we maintained the ratio at different levels: 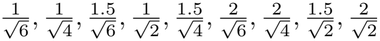, while changing the values of *µ*_1_, *µ*_2_ and *σ*. For all of the above scenarios we maintained the correlation coefficient within the family is 0.3.

The correlation between the sibs of a same sib-ship might affect the powers. To obtain the effect of correlation coefficient between the family members, we conducted simulations from N(0,2) distributions under three different correlation levels as 0, 0.3 and 0.9.

It is important to obtain the effect of family composition and size on the methods. So under the correlation level 0.3 we simulated the sib-ship sizes under a truncated Poisson distribution with 4 choices of *λ*: 1.2, 2.2, 3.2 and 4.2.

To benchmark the proposed methods against established alternatives, we simulated the powers using the classical methods like paired t-test or Wilcoxon signed-rank test under both N(0,2) and 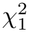 with *ρ* = 0.3. Further, to evaluate the statistical performance of LMM with REML of family effect (Wald test for fixed effect using Kenward-Roger correction [REF]), we also include this stratagy in a few simulation settings (see Supplementary Figure S1).

### 4.3 Simulation Results

Below we summarize the important results from the simulations. The detailed results are presented in Tables 1-5 and Supplementary Tables S1 and S2.

**Table 1:** Simulation results under N(0,2) distribution with different correlation coefficients.

| **Correlation Coefficients** | **0** |  | **0.3** |  | **0.9** |  |
|---|---|---|---|---|---|---|
| **Difference b/w Mean** | Parametric | Non-parametric | Parametric | Non-parametric | Parametric | Non-parametric |
| 0 | 0.049 | 0.053 | 0.053 | 0.054 | 0.049 | 0.044 |
| 1 | 0.882 | 0.663 | 0.932 | 0.814 | 1.000 | 1.000 |
| 1.5 | 0.996 | 0.939 | 1.000 | 0.986 | 1.000 | 1.000 |
| 2 | 1.000 | 0.998 | 1.000 | 1.000 | 1.000 | 1.000 |

**Table 2:**
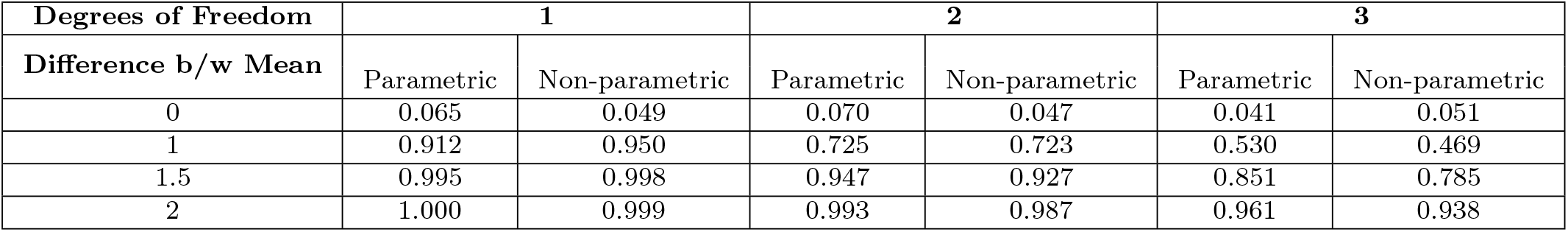
Simulation results under *χ*^2^ distribution with different degrees of freedoms with *ρ* = 0.3.

| **Degrees of Freedom** | **1** |  | **2** |  | **3** |  |
| **Difference b/w Mean** | Parametric | Non-parametric | Parametric | Non-parametric | Parametric | Non-parametric |
|---|---|---|---|---|---|---|
| 0 | 0.065 | 0.049 | 0.070 | 0.047 | 0.041 | 0.051 |
| 1 | 0.912 | 0.950 | 0.725 | 0.723 | 0.530 | 0.469 |
| 1.5 | 0.995 | 0.998 | 0.947 | 0.927 | 0.851 | 0.785 |
| 2 | 1.000 | 0.999 | 0.993 | 0.987 | 0.961 | 0.938 |

The two methods maintain the desired level of significance (corresponding to *γ* = 0) in all simulation scenarios where the data is simulated by maintaining the corresponding assumptions [Table 1-2]. Simulations under a skewed distribution like 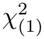 or 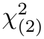 the parametric method shows inflated type 1 error. But for 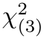 distribution, the type 1 error maintains the desired level [Figure 2]. With respect to power, we first note some findings irrespective of the distribution of the quantitative trait. Of course, the power of detecting association increases as the gap between the mean of the affected individuals and that of the unaffected ones increases [Table 1-2]. As expected, the power decreases as the variance of the quantitative trait increases and the data becomes more noisy. Obviously, if the distribution of the QT follows a normal distribution the power of the parametric test dominates that of the non-parametric one. We notice that, under N(0,2) distribution the power increases with the increment of correlation coefficient within the family members in both parametric and non-parametric setup. In fact as the correlation coefficient increases, the gaps between the powers of parametric and non-parametric setup reduces and coincides at very high value of *ρ* = 0.9 [Figure 3]. Under 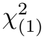 and 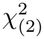 distribution, we observe that the power of the non-parametric test dominates its parametric counterpart, though the gap between them decreases with the increment of the corresponding degrees of freedoms [Figure 4]. We reduced the skewness in the 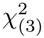 distribution to make the data comparatively symmetric around its mean and the parametric setup overpowers the non-parametric one [Figure 4]. The comparison between the classical methods and the proposed methods show that under both N(0,2) and 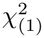, the proposed method overpowers the classical method [Figure 6]. If the distribution is same irrespective of its type, the test maintains symmetry of power [Figure 5] with respect to the equal positive and negative magnitude of *γ* [Table 3]. We also note that the power increases as the ratio 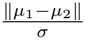 increases and the power is almost same [Table 4 - 5] if the ratio is same even after changing the values of *µ*_1_, *µ*_2_ and *σ* [Figure 7].

**Table 3:** Simulation results for symmetric mean differences under N(0,2) and 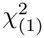 distribution with *ρ* = 0.3.

| \|**μ<sub>1</sub>** − **μ<sub>2</sub>**\| | **Positive Mean Shift** |  | **Negative Mean Shift** |  |
|---|---|---|---|---|
|  | **Parametric** | **Non-parametric** | **Parametric** | **Non-parametric** |
| Under N(0,2) |  |  |  |  |
| 1 | 0.932 | 0.814 | 0.925 | 0.823 |
| 1.5 | 1.000 | 0.986 | 0.999 | 0.987 |
| 2 | 1.000 | 1.000 | 1.000 | 1.000 |
| Under χ<sup>2</sup><sub>(1)</sub> |  |  |  |  |
| 1 | 0.912 | 0.950 | 0.942 | 0.977 |
| 1.5 | 0.995 | 0.998 | 0.998 | 1.000 |
| 2 | 1.000 | 0.999 | 1.000 | 1.000 |

**Table 4:** Simulation results with respect to the ratio 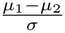 under Normal distributions with *ρ* = 0.3.

| $\frac{\mu_1-\mu_2}{\sigma}$ | **Parametric Scheme** | **Non-parametric Scheme** |
|---|---|---|
| $\frac{1}{\sqrt{6}}$ | 0.522 | 0.376 |
| $\frac{1}{\sqrt{4}}$ | 0.687 | 0.517 |
| $\frac{1.5}{\sqrt{6}}$ | 0.853 | 0.691 |
| $\frac{1}{\sqrt{2}}$ | 0.932 | 0.814 |
| $\frac{1.5}{\sqrt{4}}$ | 0.947 | 0.857 |
| $\frac{2}{\sqrt{6}}$ | 0.971 | 0.909 |
| $\frac{2}{\sqrt{4}}$ | 0.992 | 0.981 |
| $\frac{1.5}{\sqrt{2}}$ | 1.000 | 0.986 |
| $\frac{2}{\sqrt{2}}$ | 1.000 | 1.000 |

**Table 5:** Simulation results with respect to the ratio 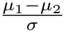 under scaled 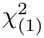 distributions with *ρ* = 0.3.

| $\frac{\mu_1-\mu_2}{\sigma}$ | **Parametric Scheme** | **Non-parametric Scheme** |
|---|---|---|
| $\frac{1}{\sqrt{6}}$ | 0.590 | 0.684 |
| $\frac{1}{\sqrt{4}}$ | 0.725 | 0.816 |
| $\frac{1.5}{\sqrt{6}}$ | 0.832 | 0.900 |
| $\frac{1}{\sqrt{2}}$ | 0.912 | 0.950 |
| $\frac{1.5}{\sqrt{4}}$ | 0.937 | 0.964 |
| $\frac{2}{\sqrt{6}}$ | 0.956 | 0.975 |
| $\frac{2}{\sqrt{4}}$ | 0.992 | 0.996 |
| $\frac{1.5}{\sqrt{2}}$ | 0.995 | 0.998 |
| $\frac{2}{\sqrt{2}}$ | 1.000 | 0.999 |

**Fig 2:**
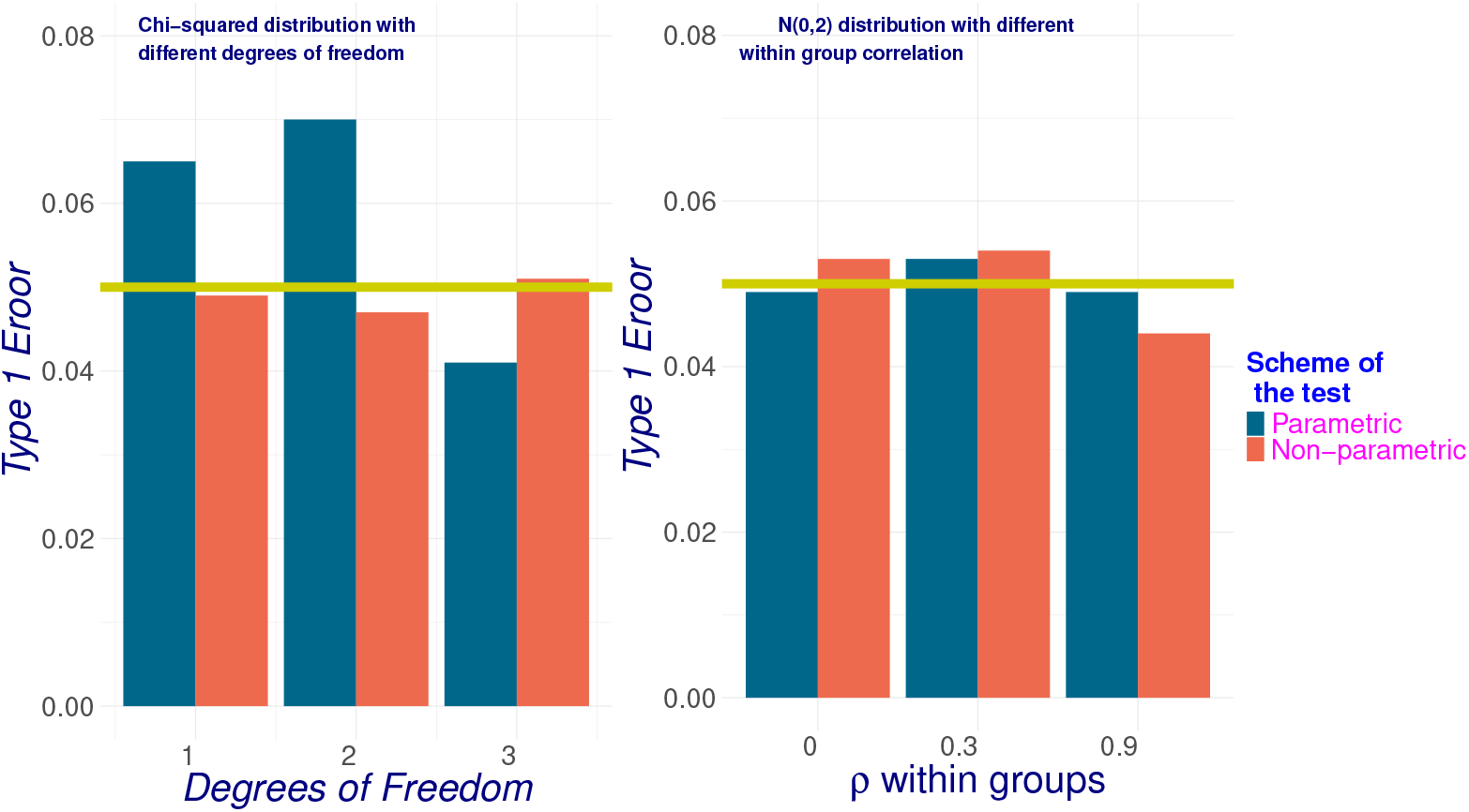
Empirical type 1 error of the proposed methods

**Fig 3:**
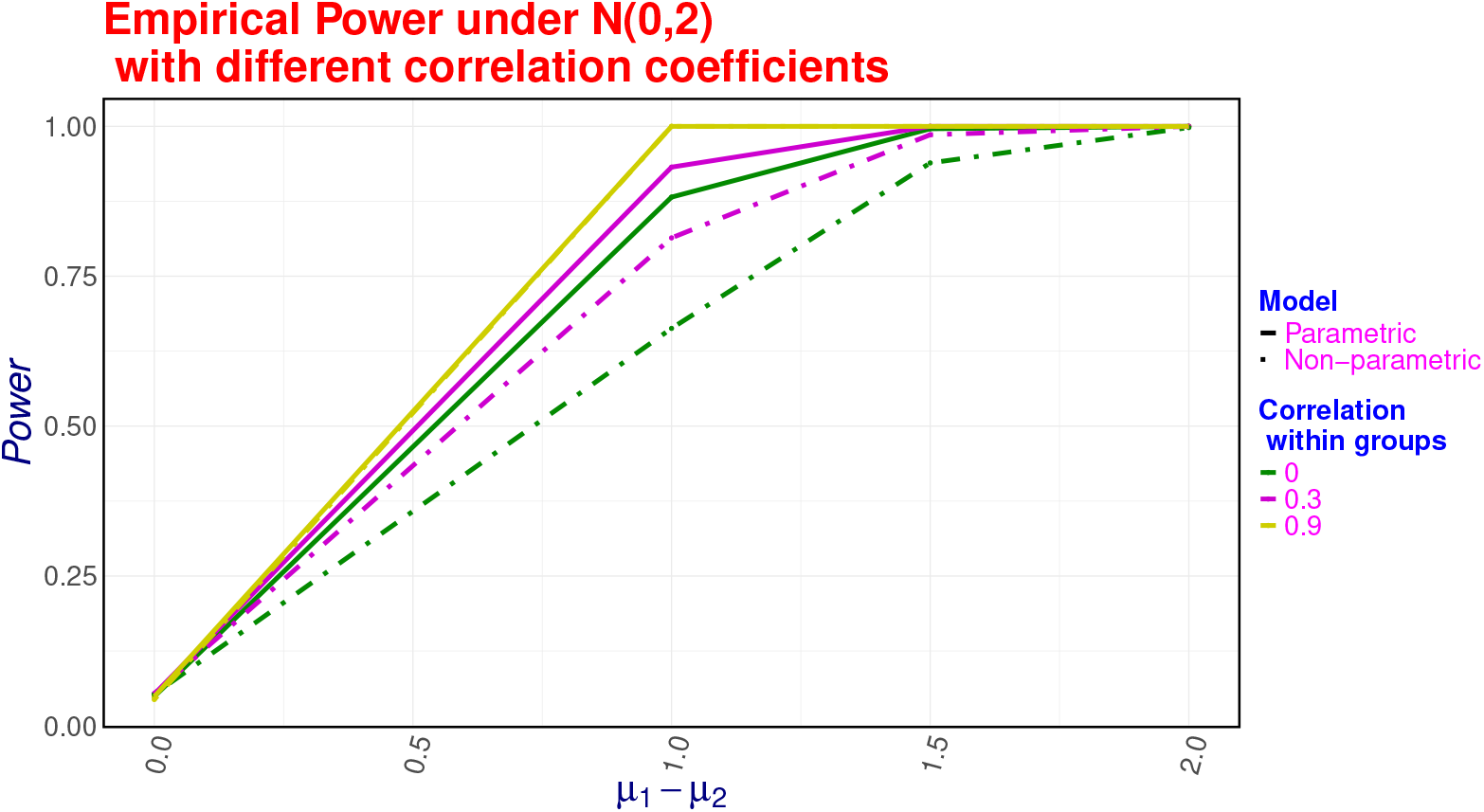
Empirical power under normal (0,2) distributions

**Fig 4:**
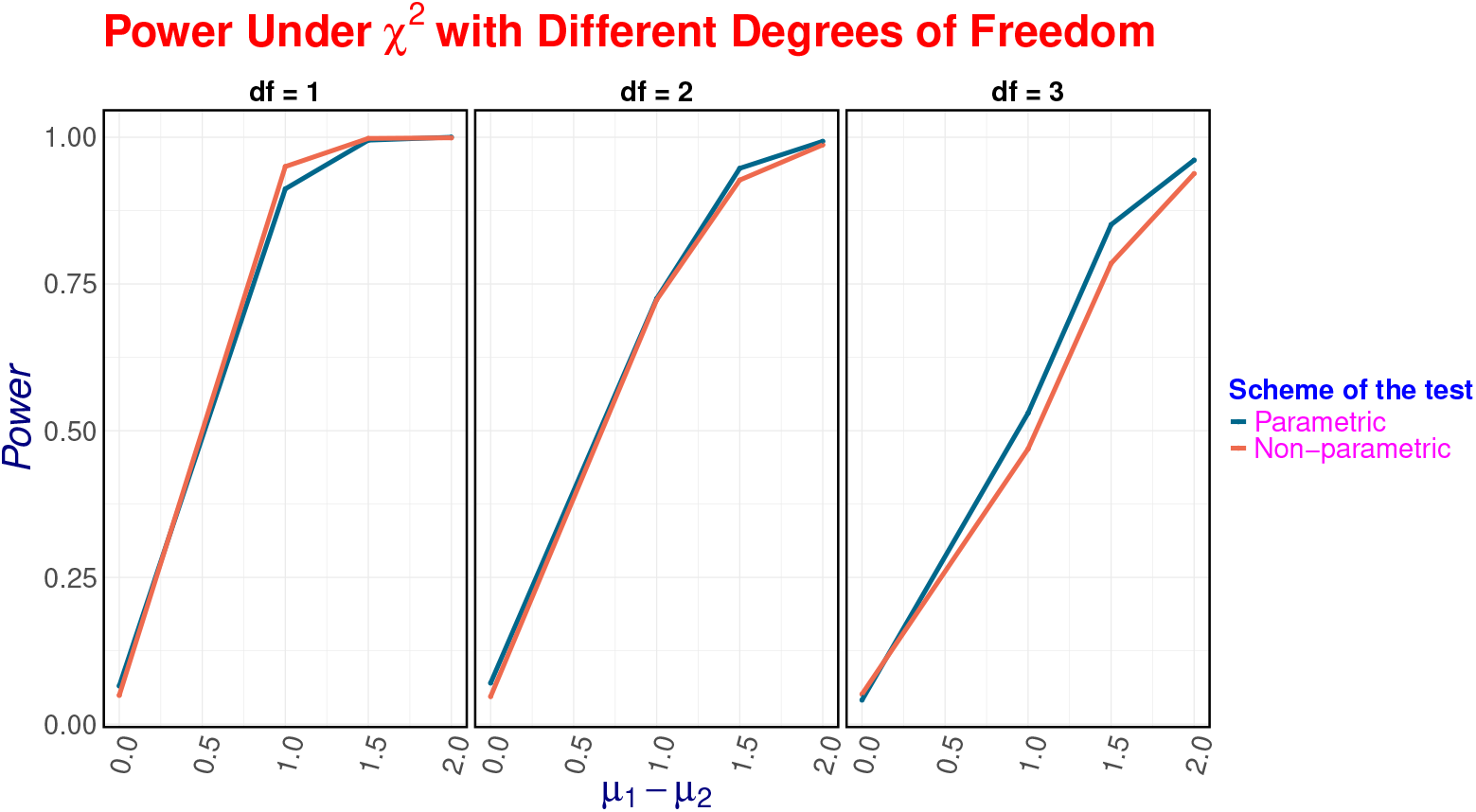
Empirical power under 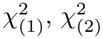 and 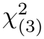 distribution with *ρ* = 0.3

**Fig 5:**
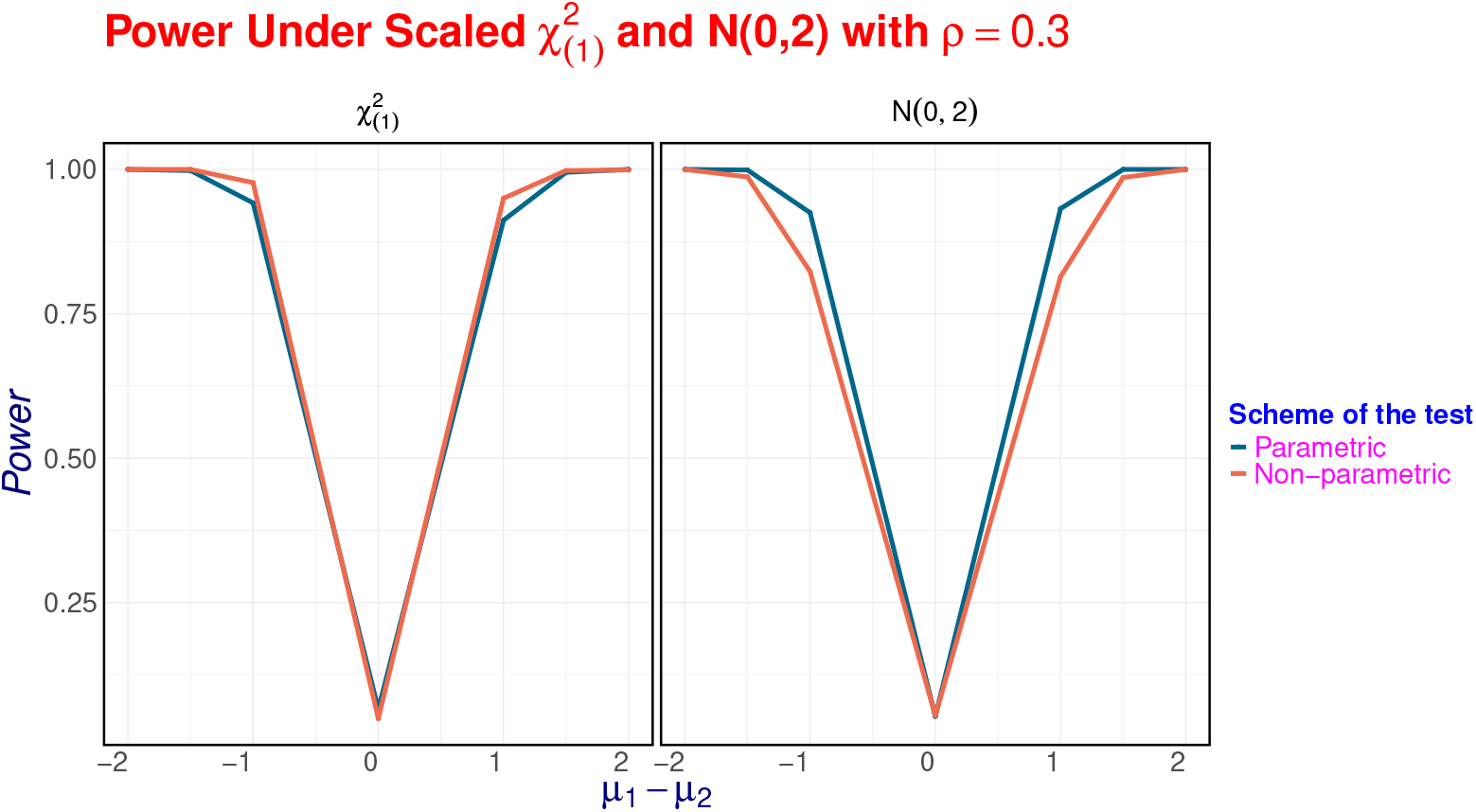
Symmetry of the tests under N(0,2) and 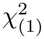 distribution with *ρ* = 0.3. It is observed that the empirical power are more or less similar at the absolute effect sizes.

**Fig 6:**
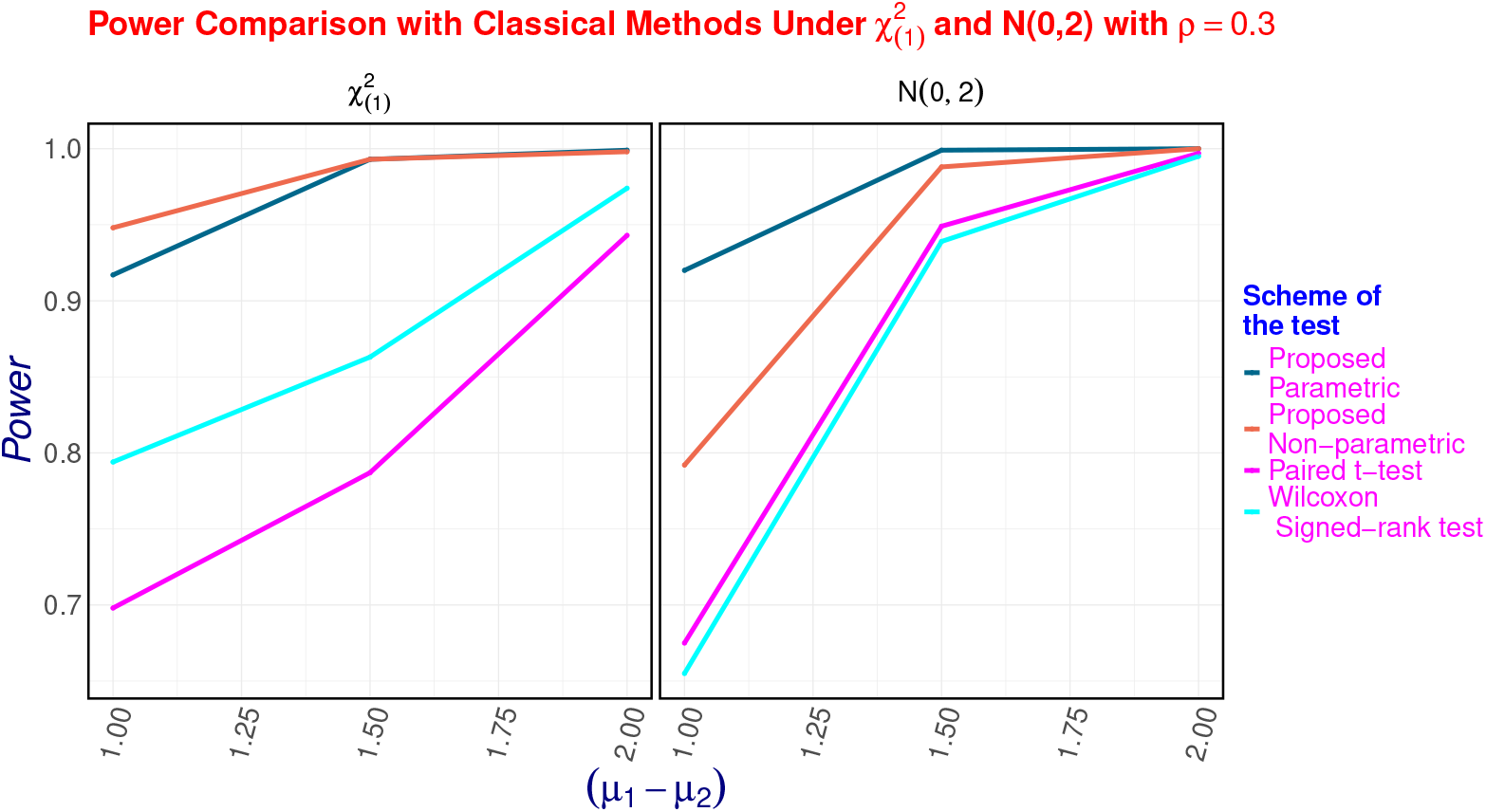
Comparison of the methods with classical methods under N(0,2) and 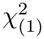 distribution with *ρ* = 0.3

**Fig 7:**
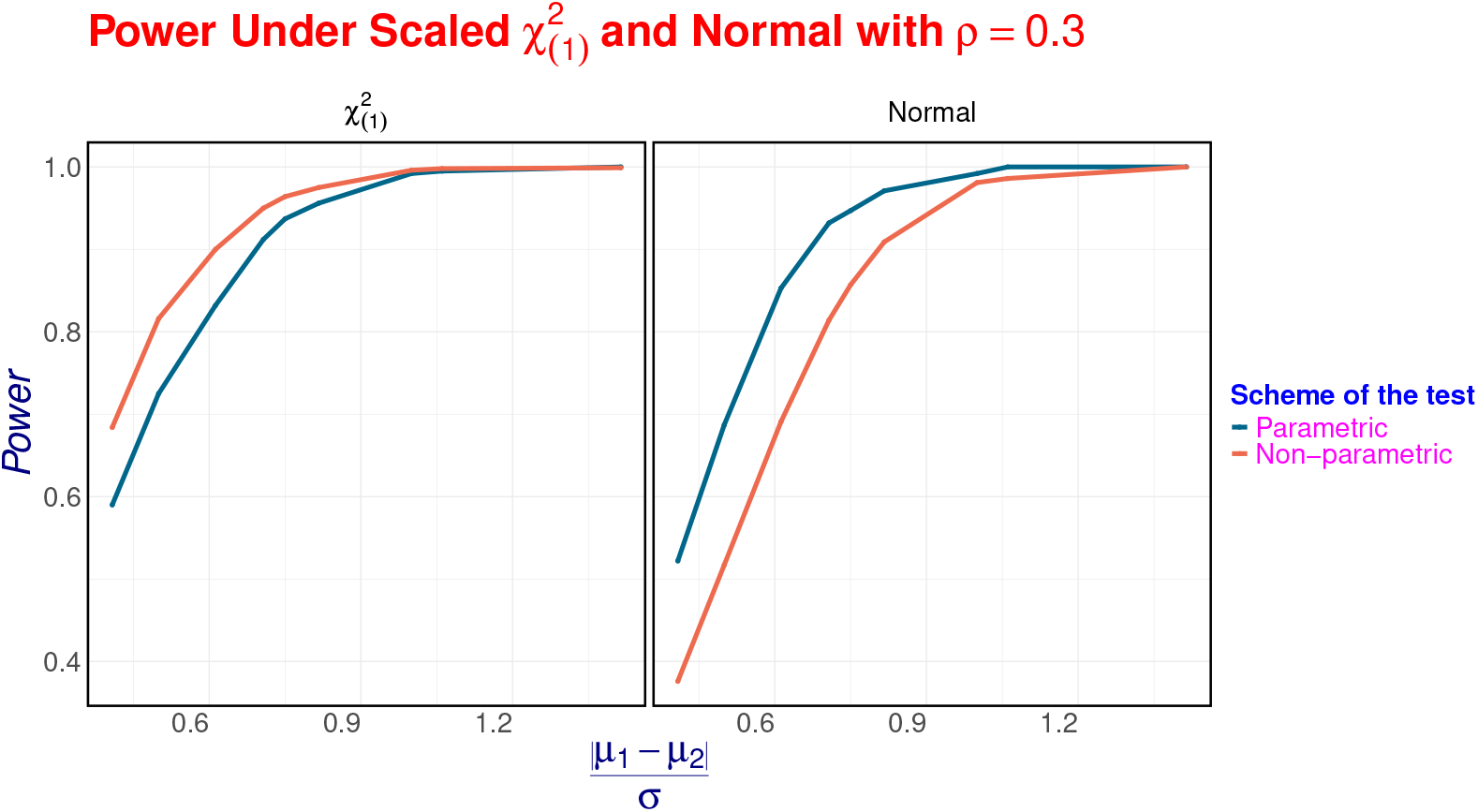
Empirical power under Normal and 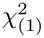 distribution with *ρ* = 0.3 wrt 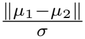

We notice that as the *λ* increases, so do the power of the tests due to increase of large sib-ship number in the data [Supplementary Table S1 - S2].

#### Robustness to Misspecified Correlation Structures

To evaluate robustness beyond the compound symmetric assumption, we generated data under two alternative within-family dependence structures: an autoregressive AR(1) model and a heterogeneous compound symmetry (HeteroCS) model. In the AR(1) setting, the correlation between the *i*-th and *j*-th siblings within a family was defined as *ρ*^|*i*−*j*|^, where *ρ* denotes the baseline intra-family correlation controlling the overall strength of dependence. In the HeteroCS setting, pairwise correlations were sampled from the *U* (0, *ρ*_*max*_) and hence the dispersion of paiwise correlation increases wrt *ρ*_*max*_, thereby inducing heterogeneity in the covariance structure. Inference was then performed using both the proposed parametric and nonparametric method as well as the linear mixed model (LMM) with random intercept assuming exchangeable correlation.

Results (Figure 8) show that the nonparametric procedure maintained Type-I error close to the nominal level across both misspecified settings. This stability arises from the permutation-based inference strategy, which relies on exchangeability of labels within families rather than explicit covariance modelling, and therefore remains less sensitive to structural misspecification. In contrast, both the parametric approach and the LMM expectedly failed to maintain Type-I error (both inflated or conservative) under misspecification of correlation structures.

**Fig 8:**
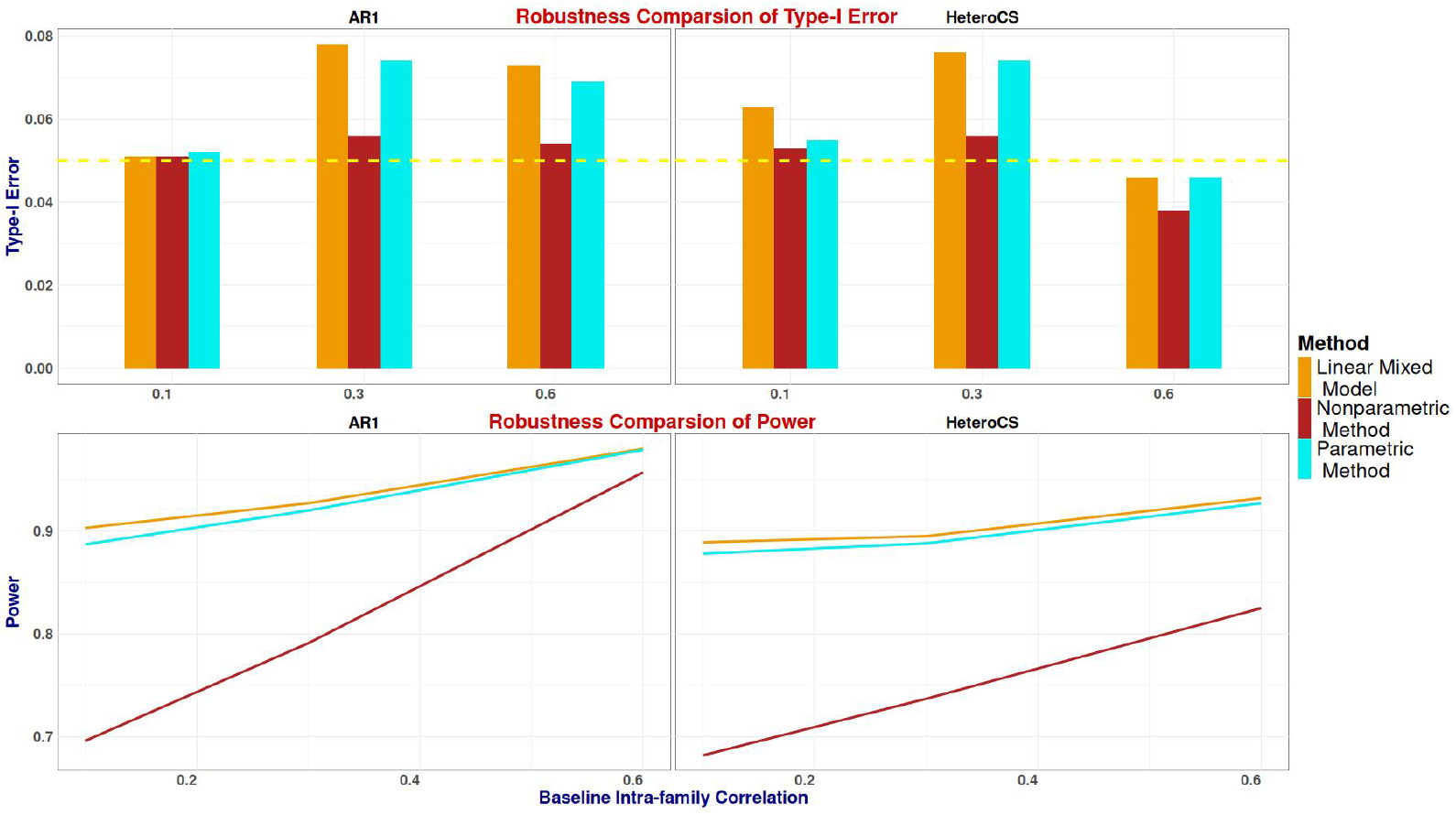
Robustness comparison under misspecified correlation structures. Data were simulated under AR(1) and heterogeneous correlation structure (HeteroCS) dependence while inference assumed exchangeable correlation. The top panels display empirical Type-I error rates and the bottom panels show power across increasing baseline intra-family correlation *ρ*. The dashed horizontal line indicates the nominal significance level *α* = 0.05.

## 5 Application to real data

### 5.1 Data description

We analysed data from the longitudinal Centre for Brain and Mind (CBM) program, previously known as the Accelerator Program for Discovery of Brain Disorders using Stem cells (ADBS)[31, 32] . The cohort comprises multi-generational families with individuals affected with major psychiatric illnesses and their unaffected first-degree relatives (FDRs), enabling the study of hereditary patterns and disease mechanisms. The families were recruited through the clinical services of National Institute of Mental Health and Neurosciences (NIMHANS), Bengaluru. For the present analyses, we included individuals with schizophrenia or bipolar disorder and their unaffected siblings. Each sibling group was independent, with no FDRs shared across different sibships. Families without unaffected siblings were excluded as they didn’t provide informative data for the planned analyses.

Participants underwent a standardized battery of neurocognitive assessments as detailed in Supplementary Methods section S1.4. The protocols used in these assessments have been described previously [33]. Sibships with missing data for any member on a given QT were excluded, resulting in variable sample sizes across traits. These QTs [Derived from the assessments mentioned in Table 6] and demographic variables (age and years of education) were used a primary phenotypes in the subsequent analyses. We also have the ‘sex’ for each patient.

**Table 6:**
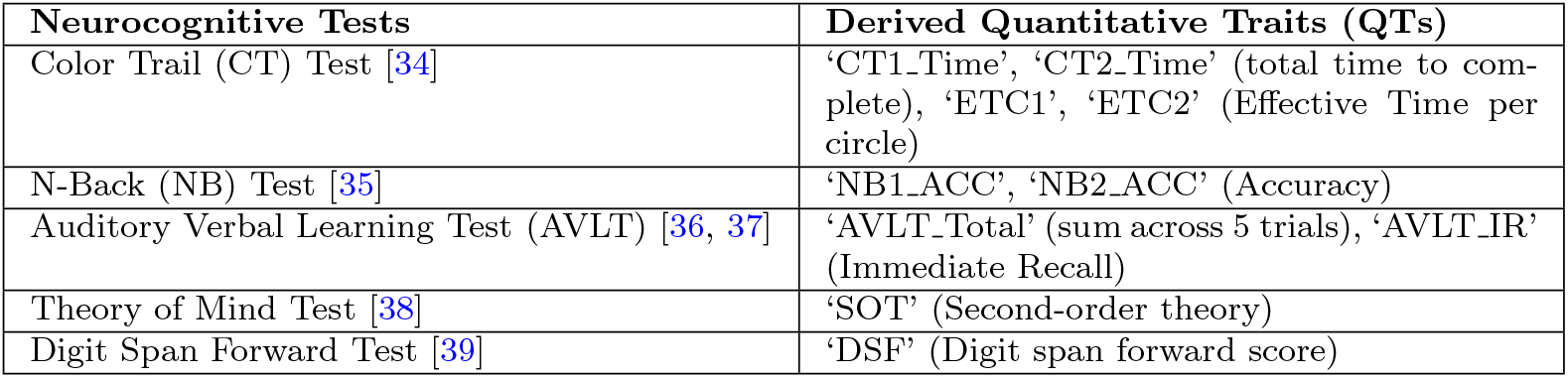
Neurocognitive assessments and derived quantitative traits (QTs)

| **Neurocognitive Tests** | **Derived Quantitative Traits (QTs)** |
|---|---|
| Color Trail (CT) Test [34] | ‘CT1_Time’, ‘CT2_Time’ (total time to com-plete), ‘ETC1’, ‘ETC2’ (Effective Time per circle) |
| N-Back (NB) Test [35] | ‘NB1_ACC’, ‘NB2_ACC’ (Accuracy) |
| Auditory Verbal Learning Test (AVLT) [36, 37] | ‘AVLT_Total’ (sum across 5 trials), ‘AVLT_IR’ (Immediate Recall) |
| Theory of Mind Test [38] | ‘SOT’ (Second-order theory) |
| Digit Span Forward Test [39] | ‘DSF’ (Digit span forward score) |

### 5.2 Analysis and results

Analyses were performed on 10 QTs for both schizophrenia and bipolar disorder, with additional covariates including age, sex and years of education. The detailed results are given in Supplementary Table S5 and Supplementary Table S6. Spplementary Table S3 summarizes the number of sibships available for each binary trait, after removal of individuals with missing data.

The majority of the quantitative traits (QTs) we examined consisted of count variables and showed a high degree of skewness. Therefore, we primarily employed the non-parametric version of the proposed test as the normality assumption would not be appropriate for most of the traits. Further, one of the relatively less skewed traits ‘AVLT Total’ was also analyzed using the parametric approach after applying the *log*(1 + *AV LT Total*) transformation. The results based on the parametric approach are shown in Supplementary Table S4. The details are mentioned in the Supplementary Methods section S1.5.

In the schizophrenia dataset(Supplementary Figure S2), 8 QTs showed correlation with disease status at a nominal level of significance (*p <* 0.05). After applying the Benjamini-Hochberg correction, all 8 QTs remained significant at a 5% FDR. Significant associations were observed for ‘CT1 Time’, ‘CT2 Time’, ‘ETC1’, ‘ETC2’, ‘NB1 Acc’, ‘AVLT Total’, ‘AVLT IR’ and ‘DSF’ [Supplementary Figure S2].

Schizophrenia was associated with lower scores in the ‘AVLT’, ‘N-Back’ and ‘Digit Span’ cluster reflecting deficits in the verbal learning, working memory and attention respectively (Supplementary Figure S3). In contrast, higher completion times in the ‘Color Trails Tests’ also indicate impairment in processing speed and cognitive flexibility. These findings are consistent with prior evidence of cognitive deficits in schizophrenia [40].

In the bipolar disorder dataset, 4 QTs were nominally significant (*p <* 0.05), all of which except ‘ETC1’ remained significant after FDR correction. Significant associations were observed for ‘CT2 Time’, ‘ETC2’ and ‘AVLT IR’ [Supplementary Figure S2]. Bipolar disorder was associated with lower values of ‘AVLT IR’ [Supplementary Figure S4] and longer time taken for color trails tests which is consistent with prior studies [41].

We identified multiple quantitative traits (QTs) significantly associated with both schizophrenia and bipolar disorder. Since analyzing by the proposed method uses family-based data, the cohort has minimal genetic and lifestyle variation between probands and their matched controls. Hence, the differences between the values of the QTs between cases and controls emphasize on the association with the binary trait with higher precision. To examine the interrelationships among the QTs identified, we calculated pairwise Spearman’s correlation coefficients and observed substantial correlations among several QTs in both disease cohorts [Supplementary Figure S6, Figure 9]. These inter-dependencies suggest the presence of shared or overlapping mechanisms underlying the observed traits.

**Fig 9:**
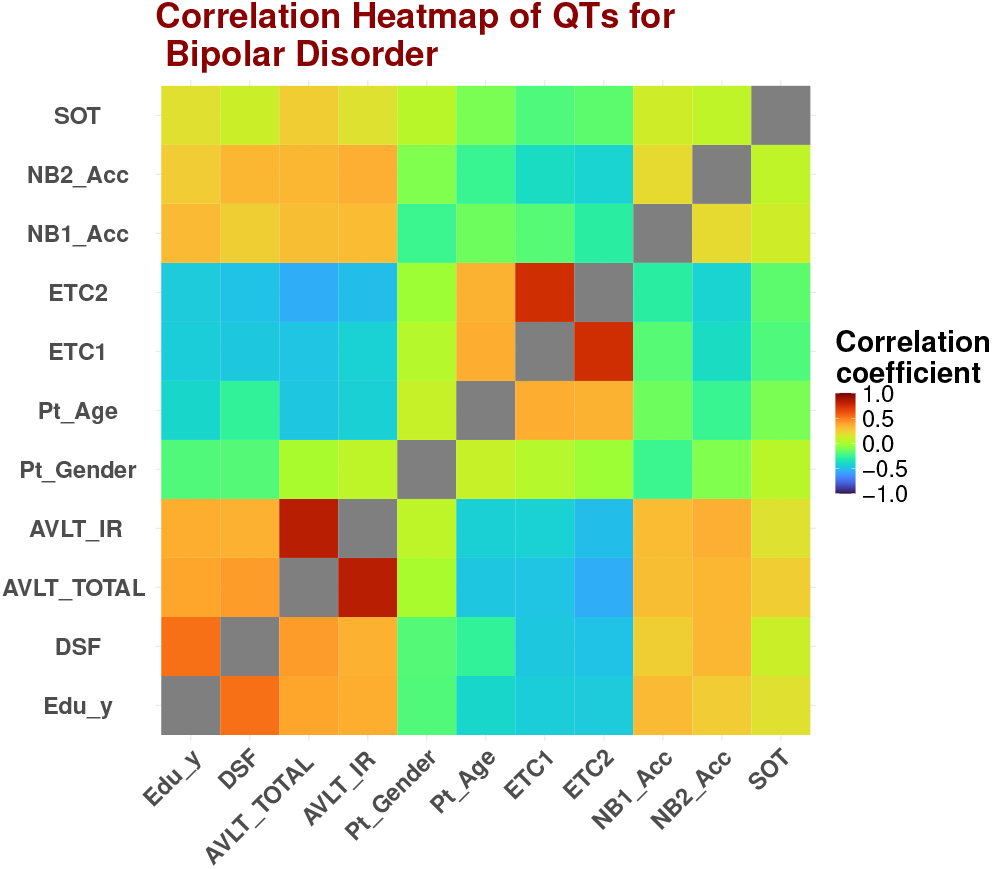
Heatmap of Correlation Coefficients among QTs for Bipolar Disorder

To accurately identify the effect of each individual QT on the disease phenotypes, it is crucial to adjust for the influence of all other QTs. Conventional approaches typically involve using covariate adjustment in a regression set up, equivalantly, obtaining residuals from regression analyses where each QT is regressed on the remaining QTs. However, due to the relatedness among individuals in the datasets and the absence of standard distributional assumptions for the QTs, we employed an in-house clustering-based adjustment method. This approach, outlined in the Supplementary Methods section S1.3.2, clusters individuals based on their covariate similarities and adjusts each QT accordingly before association analysis. Individuals are hierarchically clustered using all quantitative traits except the target trait, producing *r < n* clusters. For each cluster, the mean of the target trait is computed. Each individual’s adjusted value is obtained by subtracting their cluster-specific mean from their original. The number of clusters controls the bias–variance trade-off. We discarded ‘CT1 Time’ and ‘CT2 Time’ as covariates, since they are highly correlated with ‘ETC1’ and ‘ETC2’ (for Schizophrenia cohort Spearman’s rank correlation coefficients are 0.9964651 and 0.99443129 and for bipolar disorder 0.99729013 and 0.99604699 respectively) respectively and basically they measure the performances of the same neurocognition activities. For adjustment along with these primary QTs, we also used ‘sex’ as a covariate. Post-adjustment, we found that 2 out of the 8 significant QTs remained significantly associated with schizophrenia, and 1 out of 5 significant QTs with bipolar disorder, as summarized in Figure 10. Detailed results are presented in Table 7.

**Table 7:** Detailed Results of QTs after Adjustment for Schizophrenia and Bipolar Disorder ^*a*^.

| **Disease** | **Bipolar Disorder** |  |  |  | **Schizophrenia** |  |  |  |
|---|---|---|---|---|---|---|---|---|
| **Quantitative Trait** | **# sibships** | **Value** | **P-value** | **T<sub>obs</sub> − T<sub>μ</sub>** | **# sibships** | **Value** | **P-value** | **T<sub>obs</sub> − T<sub>μ</sub>** |
| Pt_Age | 69 | 951 | 0.814 | -20.7915 | 55 | 557.5 | 0.178 | -84.947 |
| Pt_Gender | 69 | 736 | 0.754 | -29.3915 | 55 | 770 | 0.058 | 130.5805 |
| SOT | 69 | 889 | 0.588 | -46.898 | 56 | 600 | 0.642 | 33.211 |
| Edu_y | 69 | 949.5 | 0.522 | -53.241 | 55 | 522.5 | 0.91 | -6.3475 |
| DSF | 70 | 801 | 0.428 | -75.811 | 55 | 565.5 | 0.124 | -117.2835 |
| AVLT_TOTAL | 69 | 672.5 | 0.332 | -89.399 | 55 | 241 | **0** | -314.4165 |
| ETC1 | 67 | 1082 | 0.202 | 108.9275 | 54 | 757.5 | **0** | 221.6995 |
| NB1_Acc | 70 | 671 | 0.172 | -125.8225 | 56 | 616 | 0.556 | -45.558 |
| NB2_Acc | 70 | 990.5 | 0.122 | 135.4675 | 55 | 620 | 0.748 | 24.2625 |
| AVLT_IR | 69 | 543 | 0.052 | -149.067 | 55 | 640 | 0.842 | 14.5 |
| ETC2 | 68 | 1173 | **0.004** | 248.003 | 53 | 657 | 0.364 | 66.43 |

<sup>a</sup>Bold p-values are FDR significant

**Fig 10:**
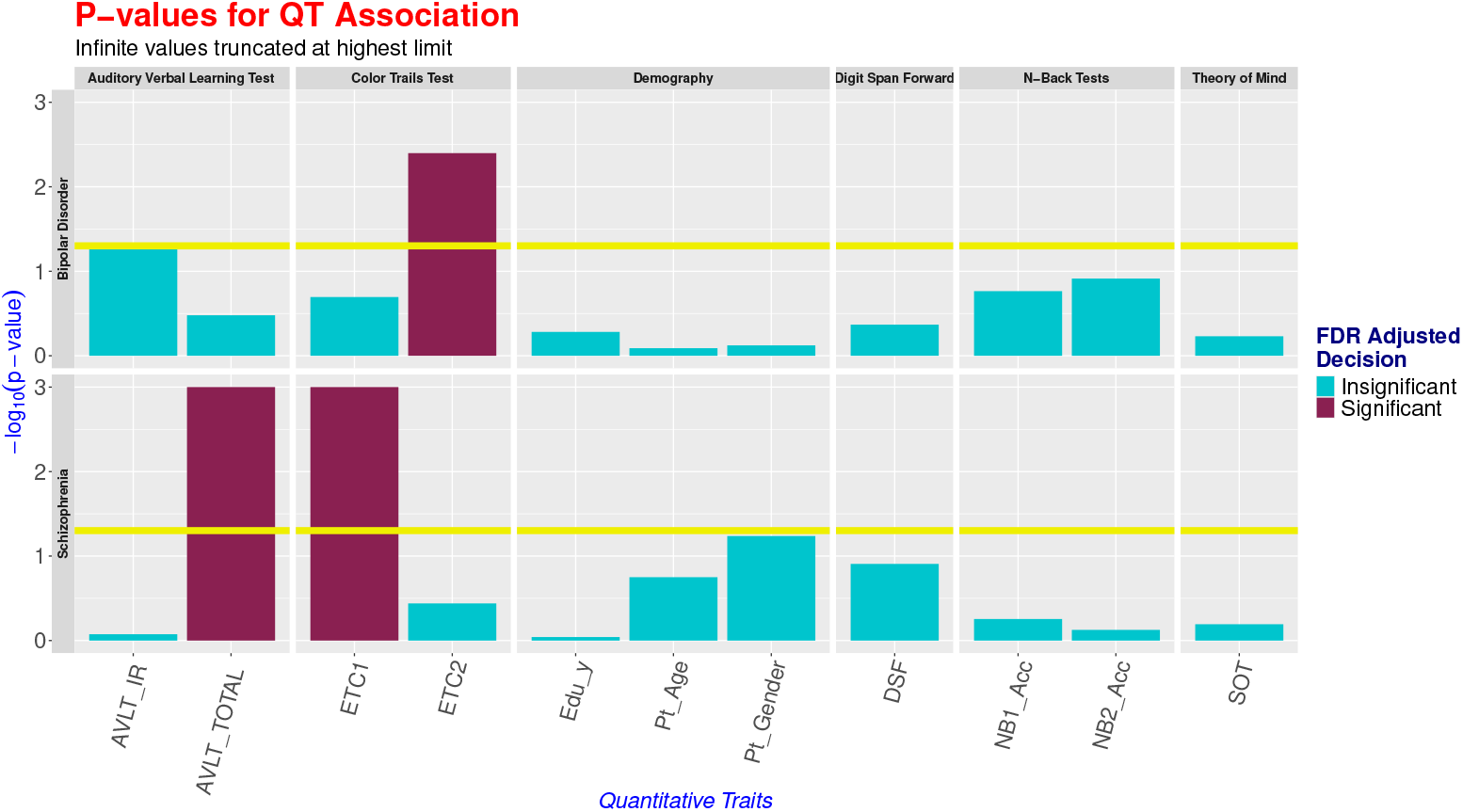
Inverted p-values of QTs after adjustment for Schizophrenia (upper-panel) and Bipolar Disorder (lower-panel). The infinite p-values are truncated at the highest limits.

The details about 95% confidence intervals of the nonparametric test statistic (*T* = (*T*_1_ + *T*_2_)) and direction of the adjusted association for schizophrenia are presented in Figure 11. Among ‘Auditory Verbal Learning Tests’ (AVLT) measures, lower values of ‘AVLT TOTAL’ and among ‘Color Trail Tests’ measures, higher values of ‘ETC1’ were significantly associated with schizophrenia after adjusting other traits as covariates. In the bipolar disorder cohort, higher values of ‘ETC2’ showed significant association after covariate adjustment.

**Fig 11:**
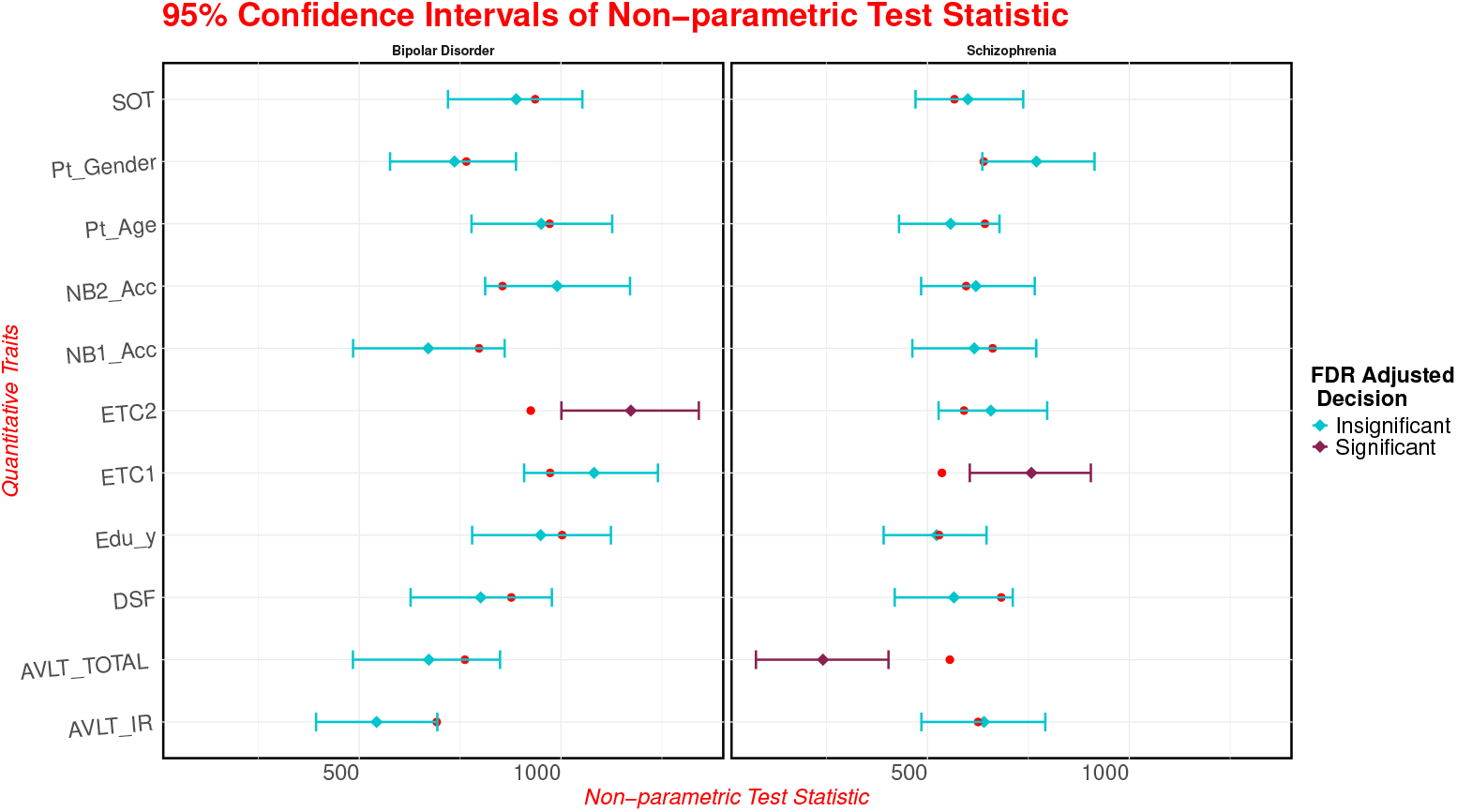
Forest plot of 95% CI of the proposed test statistic for A. schizophrenia and B. bipolar disorder (red dots are empirical null mean of the distribution)

We observed that relatively simpler cognitive tasks, such as ‘AVLT TOTAL’ and ‘ETC1’, are significantly associated with schizophrenia, whereas more complex counterparts like ‘ETC2’ show associations with bipolar disorder. This pattern is consistent with current clinical understanding and prior literature, which suggest that schizophrenia is characterized by broad, generalized cognitive impairments affecting both low- and high-level tasks [42, 43]. In contrast, bipolar disorder is typically associated with more selective deficits, particularly in domains requiring higher-order functioning. These findings imply that individuals with schizophrenia are likely to struggle even with simple cognitive tasks compared to their siblings—despite shared genetic and environmental risk factors. In contrast, bipolar patients tend to perform comparably to their siblings on basic tasks but show notable deficits in more cognitively demanding assessments.

These findings highlight a specific subset of cognitive traits with potential higher relevance to schizophrenia and bipolar disorder, adjusting for the other correlated measurements.

## 6 Discussion

We propose a parametric and a non-parametric method to identify the association between a binary phenotype and a quantitative phenotype using a mix of affected families and unrelated cases. The proposed parametric and non-parametric approaches can be thought of as extensions of the standard paired t-test and Wilcoxon signed-rank test respectively. In fact, the proposed methods address the more generic problem of equality of location parameter based on groups of correlated measurements, which can have potential applications beyond the domain of comorbidity, such as in analysis of gene-expression or other molecular quantitative trait data.

Our tests were evaluated using simulations and real data on neurocognitive assessments for individuals affected by schizophrenia/bipolar disorders and their first degree relatives. Both in the simulations and real-data assessments, the methods showed strong performance in identifying traits linked to diseases, which can help in understanding disease-risk patterns in communities. The parametric method works better when trait being studies are normally distributed, which may be a reasonable assumption for many biological markers. The non-parametric method performs better as the data increasingly deviates from normality. When the data are slightly skewed, the parametric approach generally remains optimal and maintains reliable error rates in larger datasets, making it robust for public health research. We found that the ability to detect these associations improves when the difference in the average trait values between diseased and non-diseased groups is large compared to inter-individual noise in the trait increasing consistently with the standardized effect size [44].

We observe that the increase in the correlation coefficient within family members increases the power of the tests. This is slightly counter intuitive since, higher correlation coefficient often implies a decrease in the effective sample size. But it is to be noted that, higher correlation within a family also implies that there is lesser unshared variation within a family and the change in the trait value is more due to the binary trait rather than other unaccounted factors. By using family data to match for shared risk factors like ethnicity or socioeconomic status, our methods offer a more precise way to identify traits that signal disease risk. The methods allow combining families with unrelated case-control data, making them versatile for epidemiological studies aimed at improving disease diagnosis, prevention, and targeted interventions in community health settings.

We applied our proposed methods to the Centre for Brain and Mind (CBM) cohort, comprising sibling groups enriched for schizophrenia and bipolar disorder, to evaluate associations between quantitative neurocognitive traits (QTs) and psychiatric diagnoses. The family-based design effectively controls for population stratification and shared environmental factors, enhancing the reliability of within-family comparisons. To account for interdependencies among QTs, we employed an in-house clustering-based covariate adjustment strategy. Using the non-parametric version of our test, we identified eight QT-s significantly associated with schizophrenia and five with bipolar disorder, even after controlling for multiple testing. After adjustment, two QTs remained significantly associated with each disorder, suggesting robust and independent effects. Schizophrenia showed broad associations spanning both simple (e.g., ‘AVLT Total’, ‘ETC1’) and complex tasks (e.g., ‘NB1 Acc’, ‘AVLT IR’), while bipolar disorder exhibited selective associations with higher-order cognitive traits, notably ‘ETC2’, reflecting differential cognitive impairment profiles consistent with prior clinical evidence.

While investigating a disease trait it is often quite hard to recruit healthy individuals as controls due to different reasons such as lack of incentive, misunderstanding, disbelief or false interpretation etc [45]. On the other hand, it is much easier to recruit controls from among the first degree relatives of the probands. Moreover, most (late-onset) complex diseases, sample from the parents can not be obtained. So, we focused the methodology and its assessment only on the sibship data. However, it should be possible to make minor modifications of the methods to allow parental data or other relationship types. Another potential limitation of our study is restriction to single quantitative traits. Individual analysis of the traits followed by multiple-testing adjustment (e.g. False Discovery Rate), can be used to study multiple QT-s with or without covariate adjustment, as demonstrated in our real data analysis. However, multivariate phenotype approaches accounting for cross-trait correlations may be developed in future for improvement in power and interpretations.

Onset of complex diseases is often preceded by changes in quantitative traits [46]. It is crucial to identify these precursors for better prevention, diagnosis and management. We utilize the proposed methods to identify potential precursors in the context of schizophrenia and bipolar disorder. The QT-s which we analyzed do not play a direct role in the diagnosis of the aforementioned diseases. The QT-s that show significant associations may be used as a non-invasive way to screen for individuals at risk. The methods developed here may be applicable in other settings. For example, the QT-s could be molecular features such as gene-expression from tissues and the groups of correlated measurements could come from multiple tissues or cells within the same individuals rather than from family data. However, adjusting the traits with relevant covariates (e.g. age, gender and batch effects) is important in any such application. Also, multiple testing correction would be crucial for application of these methods on large sets of omics derived traits.

## 7 Conclusion

In summary, the proposed methods offer a novel approach to datasets comprising correlated measurements, which frequently arise in statistical genetics and epidemiology. Findings from the analysis of sibling-based data on neuropsychiatric disorders support the role of select cognitive QT-s as potential endophenotypic markers and demonstrate the utility of the proposed statistical framework in extracting meaningful comorbid associations. Although the real-data demonstration is in the area of psychiatric disorders, the proposed approach is applicable to any setting where relationships between quantitative and binary traits is of interest, particular with clusters of related individuals. The strategies used here may be extendable to other contexts to enable more efficient analysis of various complex traits.

## Supplementary information

Supplementary information (methods, figures and tables) are included in a separate document along with this article.

## Declarations

### Ethics approval and consent to participate

The datasets analyzed in the real data application section were generated in an earlier study that had been approved by the relevant review boards and informed consent had been taken from all the participants. Permission to use the aforementioned data for this project was obtained from the ethics committees of both National Institute of Mental Health and Neuro Sciences (NIMHANS), Bengaluru (NIMHAN-S/EC (BEH.SC.DIV)MEETING/2024) and BRIC-National Institute of Biomedical Genomics (NIBMG), Kalyani (NIBMG/2024/1/0068).

### Consent for publication

Not Applicable

### Availability of data and materials

The real dataset used in the study can be made available upon requesting the Centre for Brain and Mind (CBM) program. Code for running the proposed methods along with synthetic dataset is available via the github repository hosted at https://github.com/sb-genome/QTRelCor.

### Author contributions

Conception and Design: Late Prof. Saurabh Ghosh, PP and SB, Provision of Neuropsychiatric Data: SJ, MP, and BV. Execution of the study: PP, Drafting of Manuscript: PP, Editing of Manuscript: SB, PP and PJ. Critical review and feedback: SB, SJ, MP, BV, SBN, PJ. Overall Supervision: SB. All authors read and approved the final manuscript.

## Acknowledgements

The cohort data came from the Centre for Brain and Mind program conducted by a team including the authors from NIMHANS, Bengaluru. The authors sincerely thank the patients and family members of the cohort. The authors would also like to thank the Human Genetics Unit of the Indian Statistical Institute (ISI), Kolkata, the BRIC National Institute of Biomedical Genomics (BRIC-NIBMG), Kalyani, the National Institute of Mental Health and Neuro Sciences (NIMHANS), Bengaluru and the Regional Centre for Biotechnology (RCB), Faridabad for providing infrastructural support to do the work. Lastly, the authors thank late Prof. Saurabh Ghosh of Human Genetics Unit of the ISI, who was involved in conceiving the project and was associated with it until his sad demise.

