## Supplementary Notes for "QT-RelCor: Statistical Inference for Comorbidity of Quantitative Traits and Disease from Related Individuals"

Supplementary Materials

### S1 Supplementary Methods

#### S1.1 Details of Comparison Study between Paired t-test and 2-sample t-test

We observe that the power of the proposed test increases with the intra-family correlation coefficient of the quantitative traits (QTs). The underlying intuition has been discussed in the main article. To better illustrate this behavior, we compare the power of the paired t-test with the two-sample t-test under identical conditions. Since our proposed parametric method generalizes the classical t-tests, and the nonparametric method serves as their distribution-free counterpart, this comparison elucidates the statistical reasoning.

Consider $Y_{i1}$ and $Y_{i2}$ as the QTs of the $i$-th family, where the first sibling is affected by the binary trait (case) and the second is unaffected (control). We assume that $(Y_{i1}, Y_{i2})$ follows a bivariate normal distribution:

$$Y_{i1} = \mu d_{i1} + e_{i1}; \forall i = 1, 2, ..., n$$
$$Y_{i2} = \mu d_{i2} + e_{i2}; \forall i = 1, 2, ..., n$$
$$d_1 = 1 \text{ and } d_2 = 0$$
$$\text{where, } e \sim N_2(\tau, \tau; \psi^2, \psi^2; \rho)$$
$$\text{Hence, } \mathbf{Y}\text{is following Bivariate Normal Distribution}$$
$$E[Y_1] = \mu + \tau$$
$$E[Y_2] = \tau$$
$$Var[Y_1] = Var[Y_2] = \psi^2$$
$$Cov(Y_1, Y_2) = \rho$$
$$\text{For paired t-test, define } D_i = (Y_{i1} - Y_{i2})$$
$$\bar{D} = \frac{1}{n}\sum_{i=1}^{n} D_i$$
$$= \frac{1}{n}\sum_{i=1}^{n} (Y_{i1} - Y_{i2})$$
$$\text{Hence, } E[D] = E[\bar{D}] = \mu, \text{ the difference between the mean of cases and controls}$$

$$Var[D] = Var[Y_1 - Y_2]$$
$$Var[D] = Var[Y_1] + Var[Y_2] - 2Cov[Y_1, Y_2]$$
$$Var[D] = 2\psi^2(1 - \rho)$$
$$\text{Hence, } Var[\bar{D}] = \frac{2\psi^2(1 - \rho)}{n}$$
$$\text{So, } Var[\bar{D}] \downarrow \rho \uparrow$$

Clearly, as the correlation coefficient $\rho$ increases, the variance of the mean difference $\bar{D}$ decreases, thereby increasing the power of the paired t-test.
This analysis confirms that the proposed method behaves in a statistically coherent manner, mirroring standard t-test properties. Moreover, it emphasizes that in case-control studies, especially when effect sizes are small, a paired testing strategy is generally more powerful and reliable than an unpaired (population-based) approach.

#### S1.2 Relation to Generalized Least Squares (GLS) and Linear Mixed Models

##### S1.2.1 Relation to GLS

The proposed parametric estimator would be equivalent to a Generalized Least Squares (GLS) estimator if the variance-covariance matrix is fully specified. When the matrix is unknown, the GLS computation is done at each iteration within the current estimate of the correlation matrix. Let $\mathbf{X}$ denote the vector of quantitative trait values and $\mathbf{D}$ the binary disease indicator. The model can be written as

$$\mathbf{X} = \mathbf{Y}\boldsymbol{\mu} + \boldsymbol{\epsilon}, \qquad \boldsymbol{\epsilon} \sim N(\mathbf{0}, \Sigma(\theta)), \quad \text{where } \mathbf{Y} = [\mathbf{D} \ \ (1 - \mathbf{D})], \ \boldsymbol{\mu} = (\mu_1, \mu_2)^\top.$$

Under compound symmetry,

$$\Sigma_k = \sigma^2\{(1-\rho)I_{m_k} + \rho J_{m_k}\} = \sigma^2 R_k,$$

Since different families are assumed independent, the full covariance matrix $\Sigma$ is block-diagonal with blocks $\Sigma_k$, i.e. $\Sigma = \text{diag}(\Sigma_1, \ldots, \Sigma_n) = \sigma^2 R$, where $R$ is the correlation matrix of the QT data. Hence, the GLS estimator takes the form

$$\hat{\boldsymbol{\mu}}_{GLS} = (\mathbf{Y}^\top R^{-1}\mathbf{Y})^{-1}\mathbf{Y}^\top R^{-1}\mathbf{X}.$$

and the corresponding variance estimator is,

$$\hat{\sigma}^2 = \frac{1}{n}(X - Y\hat{\mu}_{GLS})^T R^{-1}(X - Y\hat{\mu}_{GLS})$$

where $n = \sum_k m_k$, is the total number of individuals $\rho$ is estimated using its profile likelihood based on the current $\hat{\mu}_{GLS}$ and $\hat{\sigma}^2$

In the present formulation the inverse correlation matrix is obtained analytically, where the diagonal and off–diagonal elements correspond to $C_k(\rho)$ and $h_k(\rho)$ respectively. Consequently, maximization of the likelihood yields estimating equations identical to GLS, demonstrating that the custom ML estimator recovers the GLS solution without repeated numerical inversion of $\Sigma$.

Under compound symmetry the correlation block can be written as

$$R_k = (1-\rho)I_{m_k} + \rho J_{m_k}, \qquad J_{m_k} = \mathbf{1}\mathbf{1}^\top.$$

Applying the Sherman–Morrison identity to matrices of the form $(I+\alpha J)^{-1}$ yields

$$R_k^{-1} = \frac{1}{1-\rho}\left(I_{m_k} - \frac{\rho}{1+(m_k-1)\rho}J_{m_k}\right).$$

Since the diagonal elements of $I_{m_k}$ and $J_{m_k}$ equal one and the off-diagonal elements of $J_{m_k}$ equal one, the diagonal and off-diagonal entries of $R_k^{-1}$ become

$$C_k = \frac{\rho(2-m_k)-1}{(\rho-1)(m_k\rho+1-\rho)}, \qquad h_k = -\frac{\rho}{(\rho-1)(m_k\rho+1-\rho)}.$$

These coincide with the coefficients appearing in the likelihood, demonstrating that the proposed ML formulation corresponds to an analytic inversion of the compound-symmetric covariance matrix.

##### S1.2.2 Comparison with LMM using REML

The proposed parametric model can be viewed as a linear mixed model (LMM) with a random intercept for each family. The covariance structure admits the representation

$$X_{ik} = \mu D_{ik} + b_k + \varepsilon_{ik},$$

where $b_k \sim N(0,\tau^2)$ is the family specific random effect and $\varepsilon_{ik} \sim N(0,\omega^2)$ is the residual. Under this parameterization the marginal variance of an observation is

$$\mathrm{Var}(X_{ik}) = \mathrm{Var}(b_k+\varepsilon_{ik}) = \tau^2+\omega^2.$$

For two distinct members $i \neq j$ within the $k$-th family,

$$\mathrm{Cov}(X_{ik},X_{jk}) = \mathrm{Cov}(b_k+\varepsilon_{ik},\, b_k+\varepsilon_{jk}) = \mathrm{Var}(b_k) = \tau^2,$$

since the residuals are independent.

Therefore, the intra-family correlation coefficient (ICC) is

$$\rho = \frac{\mathrm{Cov}(X_{ik},X_{jk})}{\mathrm{Var}(X_{ik})} = \frac{\tau^2}{\tau^2+\omega^2}.$$

Thus, the ML estimate $\hat{\rho}$ obtained from our likelihood framework aligned closely as expected with $\frac{\hat{\tau}^2}{\hat{\tau}^2+\hat{\omega}^2}$ from the fitted LMM. The iterative ML estimation approach is more general than the LMM estimation with random intercept, as the later is only applicable in the compound symmetry situation.

We perform limited simulations to benchmark the proposed method against LMM using REML estimate. For each simulated dataset we compared (i) the proposed parametric likelihood-ratio test, (ii) the nonparametric procedure, and (iii) the LMM using Wald-type inference with Kenward-Roger (KR) correction on the disease-status as fixed effect (standard LRT or Wald Test cannot be used for testing the fixed effect obtained using LMM with REML estimate). Since fixed-effect estimation in the LMM corresponds to a GLS estimator, similar power between the parametric approach and the LMM under Gaussian settings is theoretically expected and was observed empirically (Supplementary Figure S1).

Under Normal $(0, 2)$ distribution, the proposed parametric method and the LMM exhibited comparable power across all correlation levels. This agreement is consistent with the fact that both procedures effectively estimate the same GLS solution under the correct model specification.

The Type-I error remained close to the nominal level for both approaches, indicating an appropriate calibration when the parametric assumptions hold. Under the skewed distribution $\chi^2_{(1)}$, both the parametric method and the LMM exhibited Type-I error inflation relative to the nominal level, consistent with sensitivity to deviations from normality, whereas the nonparametric approach maintained stable Type-I error control. Moreover, the nonparametric procedure achieved higher empirical power, suggesting increased robustness when distributional assumptions are violated.

#### S1.3 Covariate Adjustment Teqchniques

##### S1.3.1 Covariate Adjustment in the Parametric Framework

The parametric likelihood framework extends naturally to incorporate additional covariates. Suppose $W_{ik}$ denotes an observed covariate (e.g., age or sex) for the $i$-th individual in the $k$-th family. The linear model can then be written as

$$X_{ik} = \theta + \mu D_{ik} + \gamma W_{ik} + \varepsilon_{ik},$$

where $\mu$ represents the effect of the binary disease indicator, $\gamma$ is the covariate effect, and the error term $\boldsymbol{\varepsilon}_k \sim N(\mathbf{0}, \Sigma_k)$ with compound-symmetric covariance

$$\Sigma_k = \sigma^2\{(1-\rho)I_{m_k} + \rho J_{m_k}\}.$$

Stacking observations across families, the model admits the matrix form

$$\mathbf{X} = \mathbf{Z}\boldsymbol{\beta} + \boldsymbol{\varepsilon},$$

where

$$\boldsymbol{\beta} = (\theta, \mu, \gamma)^\top, \qquad \mathbf{Z} = \begin{bmatrix} 1 & D_{11} & W_{11} \\ \vdots & \vdots & \vdots \\ 1 & D_{ik} & W_{ik} \end{bmatrix}.$$

Conditional on $(\sigma^2, \rho)$, the maximum likelihood estimator of $\boldsymbol{\beta}$ coincides with the generalized least squares (GLS) solution

$$\hat{\boldsymbol{\beta}_{GLS}} = (\mathbf{Z}^\top R^{-1}\mathbf{Z})^{-1}\mathbf{Z}^\top R^{-1}\mathbf{X}.$$

Given $\hat{\boldsymbol{\beta}}$, the variance component $\sigma^2$ is estimated from the corresponding quadratic form, and the correlation parameter $\rho$ is obtained by solving the profile score equation

$\partial\ell/\partial\rho = 0$, analogous to the intercept-only case described in the 'Methods' section. These updates are iterated until convergence of the profile likelihood.

##### S1.3.2 Nonparametric Covariate Adjustment of Quantitative Traits

To obtain covariate adjusted association of the QT-s, residuals of each trait can be derived to remove the part explained by the other QT-s. Generally, regression-based techniques are applied to adjust covariates for independent individuals where the trait distribution is known (e.g. linear and logistic regression). For the analysis of sib-ship data with various highly skewed traits including ordinal traits, we were unable to apply such methods. Hence, the following clustering-based approach was used to derive adjusted residuals for each QT.

Suppose $X_1$ is the QT of interest and there are total p QTs, say $X_1, X_2, ..., X_p$ for n individuals. Let $X_1$ be the QT of interest. Now, we do a hierarchical clustering on the individuals based on the distance matrix of all other QTs i.e., on $X_2, X_3, ..., Xp$. Say we have r($<< n$) clusters of individuals, say $C_1, C_2, ..., C_r$. Now, we compute the mean of $X_1$ for all the r clusters $\overline{X_{C_1}}, \overline{X_{C_2}}, ..., \overline{X_{C_r}}$. Since each individual belongs to either of the r clusters, we then subtract the corresponding cluster mean from the original QT value to get the adjusted QT value, that is, $X^*_{1i} = X_{1i} - \overline{X_{C_j}}$ if $i \in C_j \forall i = 1, 2, .., n$ and $j = 1, 2, ..., r$.
As with all clustering algorithms, it is unclear how the number of clusters should be chosen. A higher number of clusters reduces the bias, but increases the variability due to smaller sample size within clusters. Conversely, too few clusters will lead to inadequate adjustment and biased effect estimation. For this analyses, we used $r = 13$, which corresponds to approximately 15% and 10% of the total number of individuals for schizophrenia and bipolar disorder respectively.

#### S1.4 Details of Neurocognitive Assessments

The types of the administered tests, for which we analyzed comorbidity, were as follows:

- Color Trails (CT) tests, consisting of Tests A and B, which measure processing speed, sustained attention, and cognitive flexibility. Scores are derived from total time taken and number of errors, yielding a metric called "effective time-per-circle."
- Auditory N-back (1-back and 2-back) tests, assessing verbal working memory by presenting consonants randomly. Scores focus on accuracy.
- Rey's Auditory Verbal Learning Test (RAVLT), evaluating verbal learning and memory by presenting and recalling sets of words. Outcome measures include RAVLT Total Score and RAVLT Immediate Recall.
- Digit Span Forward test, assessing short verbal memory by recalling sequences of digits in the same order. Performance relies primarily on the integrity of the left hemisphere.
- Theory of Mind (ToM) tests, designed to measure the understanding and interpretation of mental states of oneself and others, involving tasks like false

belief scenarios and recognizing emotions from facial expressions or understanding sarcasm.

##### S1.5 Analysis of 'AVLT_TOTAL' for Schizophrenia under Parametric Setup

To benchmark the performance of the novel likelihood ratio test (LRT)-based method, we analyzed the quantitative trait (QT) AVLT TOTAL for schizophrenia. Although this is a count variable, we applied a $\log(1+\text{AVLT TOTAL})$ transformation to achieve approximate symmetry around the mean, as illustrated in Figure S7. The transformed variable was then analyzed using the parametric approach. The QT was found to be statistically significant at the $\alpha = 0.05$ level.

Given that AVLT TOTAL is correlated with other QTs, we adjusted its values via linear regression using the remaining QTs as covariates. To ensure independence among samples for regression analysis, we included only unrelated individuals: specifically, uncorrelated probands and a single control per family. Based on the regression model, we computed the expected values of AVLT TOTAL and derived residuals by subtracting these expectations from the observed transformed values.

The parametric method was subsequently applied to the residuals. In this case, the test result was no longer statistically significant, yielding a p-value of 0.9531. This outcome suggests that adjusting AVLT TOTAL for other QTs may substantially reduce the observed signal, possibly indicating that the initial significance was partially driven by correlation with these covariates.

##### S1.6 Computational Costs and Convergence

The iterative profile likelihood estimation takes only a few seconds on a standard desktop computer for family and sample sizes as mentioned in our simulation. In case of boundary situations like too few related individuals with very small correlation, the estimation of $\rho$ may be unstable and hence, in that case dropping related individuals can provide an alternative practical option.

For the nonparametric approach, the total runtime is also a few seconds on a standard desktop for the same scenarios. There is no convergence issue for this approach since, there are no iterative solver procedure or maximization involved. Runtime for the nonparametric approach depends on the number of permutations. To increase the accuracy of $p$-values more permutations are needed at the cost of higher runtime.

### S2 Supplementary Figures and Tables

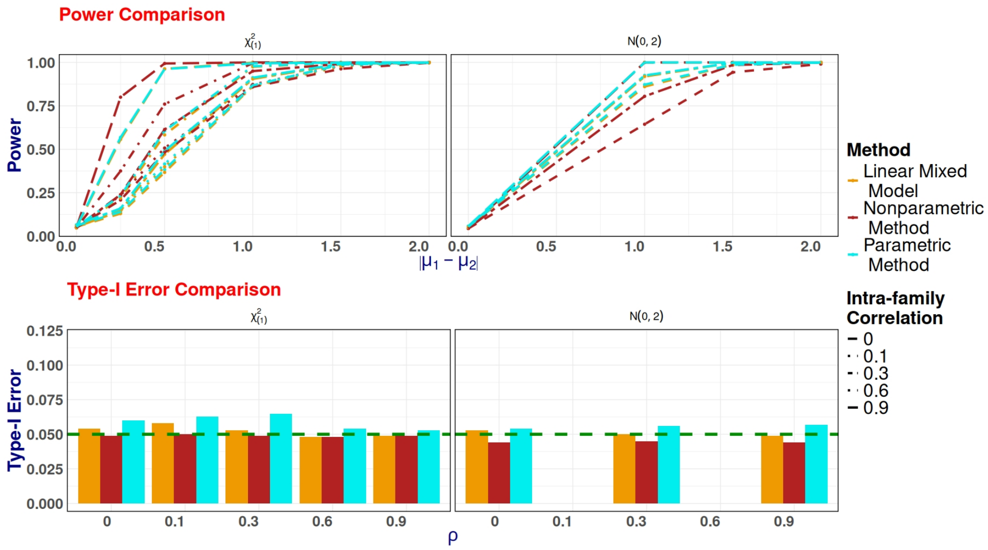


**Fig. S1**: Type-I error and Power Comparison under with LMM. All approaches remain close to the nominal $\alpha = 0.05$ level under N(0,2) but not under $\chi^2_{(1)}$ distribution except for the nonparametric approach, indicating appropriate calibration when model assumptions hold.

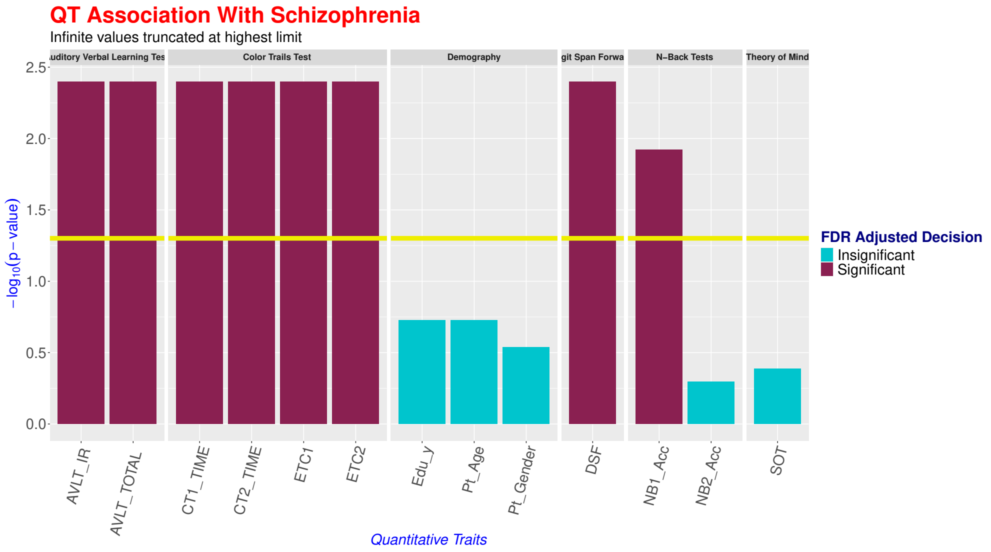


**Fig. S2**: Plot of $-\log_{10}(p - values)$ of unadjusted QTs for Schizophrenia

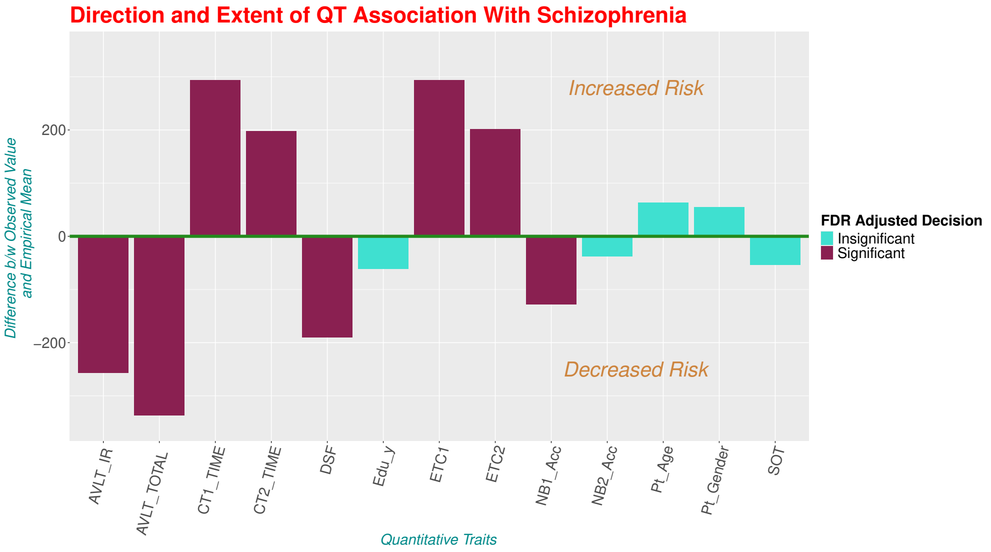


**Fig. S3**: Extent and Direction of the associated unadjusted QTs for Schizophrenia

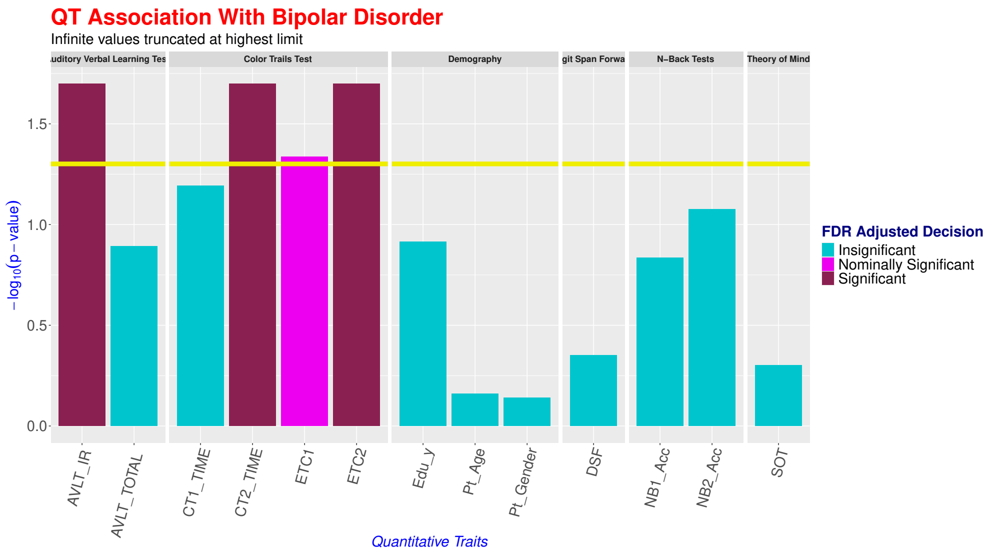


**Fig. S4**: Plot of $-\log_{10}(p - values)$ for Bipolar Disorder

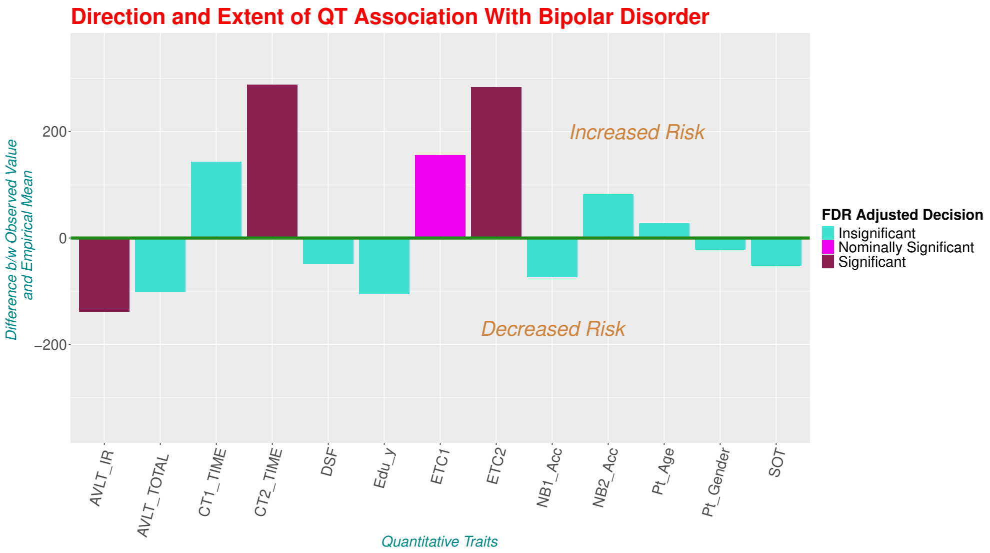


**Fig. S5**: Extent and Direction of the associated QTs for Bipolar Disorder

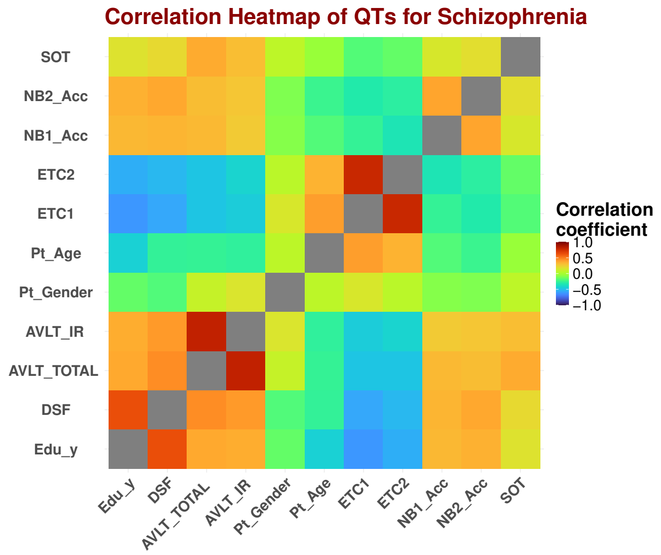


**Fig. S6**: Heatmap of Correlation Coefficients among unadjusted QTs for Schizophrenia

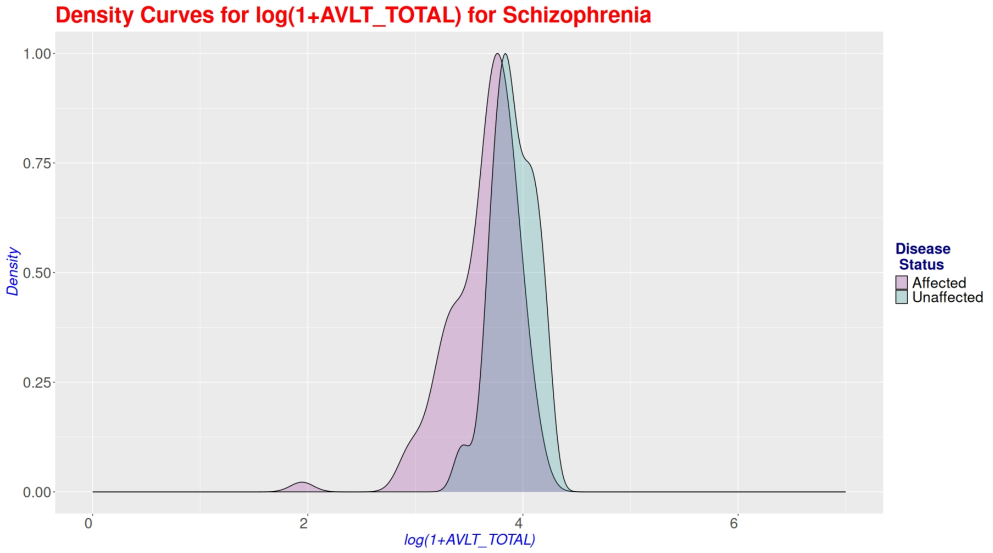


**Fig. S7**: Distribution of 'AVLT TOTAL' for Schizophrenia

**Table S1**: Simulation results with respect to $\lambda$ under Normal(0,2) distribution with $\rho = 0.3$.

| | $\gamma = 1$ | | $\gamma = 2$ | |
|---|---|---|---|---|
| $\lambda$ | Parametric | Non-parametric | Parametric | Non-parametric |
| 1.2 | 0.932 | 0.814 | 1.000 | 1.000 |
| 2.2 | 0.980 | 0.952 | 1.000 | 1.000 |
| 3.2 | 0.994 | 0.980 | 1.000 | 1.000 |
| 4.2 | 1.000 | 0.993 | 1.000 | 1.000 |

**Table S2**: Simulation results with respect to $\lambda$ under $\chi^2_{(1)}$ distribution with $\rho = 0.3$.

| | $\gamma = 1$ | | $\gamma = 2$ | |
|---|---|---|---|---|
| $\lambda$ | Parametric | Non-parametric | Parametric | Non-parametric |
| 1.2 | 0.912 | 0.950 | 1.000 | 0.999 |
| 2.2 | 0.975 | 0.989 | 0.992 | 0.999 |
| 3.2 | 0.997 | 1.000 | 1.000 | 1.000 |
| 4.2 | 0.997 | 1.000 | 1.000 | 1.000 |

**Table S3**: Summary of number of effective sib-ship

| **Name of the Binary Trait** | **Minimum** | **Maximum** | **Median** |
|---|---|---|---|
| Schizophrenia | 53 | 56 | 55 |
| Bipolar Disorder | 67 | 70 | 69 |

**Table S4**: Results for 'AVLT TOTAL' for Schizophrenia under Parametric Setup

| **Type** | **QT** | **# Sibship** | **Likelihood Ratio** | **P-value** |
|---|---|---|---|---|
| Unadjusted | AVLT_TOTAL | 55 | 32.63706 | **1.11076e-08** |
| Adjusted | AVLT_TOTAL | 55 | 33.47043 | **7.2355e-09** |

**Table S5**: Detailed results of the unadjusted QTs wrt schizophrenia

| Quantitative Trait | Number of Effective Sibship | Observed test statistic | P-value | Observed Value - Empirical Mean |
|---|---|---|---|---|
| NB2_Acc | 55 | 544 | 0.504 | -37.76199 |
| SOT | 56 | 473.5 | 0.41 | -53.84401 |
| Pt_Gender | 55 | 562 | 0.29 | 55.0285 |
| Edu_y | 55 | 374.5 | 0.188 | -61.1435 |
| Pt_Age | 55 | 867 | 0.188 | 63.038 |
| NB1_Acc | 56 | 489.5 | 0.012 | -128.2255 |
| DSF | 55 | 407 | 0.004 | -189.844 |
| CT1_TIME | 54 | 1014.5 | 0 | 293.6335 |
| CT2_TIME | 53 | 896 | 0 | 197.457 |
| AVLT_TOTAL | 55 | 208.5 | 0 | -337.085 |
| AVLT_IR | 55 | 240.5 | 0 | -256.942 |
| ETC1 | 54 | 1016 | 0 | 294.1205 |
| ETC2 | 53 | 897 | 0 | 201.271 |

**Table S6**: Detailed results of the unadjusted QTs wrt bipolar disorder

| Quantitative Trait | Number of Effective Sibship | Observed test statistic | P-value | Observed Value - Empirical Mean |
|---|---|---|---|---|
| Pt_Gender | 69 | 659 | 0.726 | -22.042 |
| Pt_Age | 69 | 1094.5 | 0.692 | 27.39249 |
| SOT | 69 | 715.5 | 0.498 | -51.76701 |
| DSF | 70 | 677 | 0.446 | -49.2365 |
| NB1_Acc | 70 | 555 | 0.146 | -73.58 |
| AVLT_TOTAL | 69 | 581.5 | 0.128 | -101.5795 |
| Edu_y | 69 | 695 | 0.122 | -105.913 |
| NB2_Acc | 70 | 771 | 0.084 | 82.668 |
| CT1_TIME | 67 | 1141 | 0.064 | 143.572 |
| ETC1 | 67 | 1163 | 0.046 | 155.248 |
| AVLT_IR | 69 | 435.5 | 0.02 | -138.173 |
| CT2_TIME | 68 | 1274.5 | 0 | 288.3005 |
| ETC2 | 68 | 1276 | 0 | 283.6775 |
